# Development of Long-Term Human Adipocyte Organoids Manifesting Aging in Response to Intermittent Hypoxia

**DOI:** 10.1101/2025.11.28.691223

**Authors:** Yunzhou Dong, Jingjing Wang, Feifei Feng, Isra H. Ali, Shahid Karim, Joshua Bock, Virend K. Somers

## Abstract

Obstructive sleep apnea (OSA) (and consequent intermittent hypoxia (IH)) is increasingly recognized as a driver of adipose tissue dysfunction, insulin resistance, and aging. However, current *in vitro* experimental models inadequately capture the long-term effects of IH on human adipocytes. Here, we report the development and optimization of a robust long-term human adipocyte organoid culture system that faithfully recapitulates IH-induced adipocyte aging *in vitro*. Human stromal vascular fraction (SVF) cells, isolated from subcutaneous abdominal fat biopsies, were embedded in Matrigel and seeded into Biofloat U-bottom 96-well plates. Using a 1:1 Matrigel-cell mixture and optimized seeding volumes (5–20 µL), we established adipocyte organoids that formed within 10–12 days and remained viable with stable morphology for up to 90 days or more. Matrigel was essential for organoid integrity, while alternative matrices such as gelatin and low-melting agarose failed to support proper organoid formation. Subcutaneous preadipocyte medium with 10% FBS from ZenBio was superior to “Advanced/F12K” medium for adipogenic differentiation and long-term maintenance.

To model OSA-related hypoxic stress, we exposed organoids to intermittent hypoxia using a programmable hypoxia chamber. IH treatment suppressed adipogenesis, as shown by reduced lipid accumulation, downregulation of adipogenic markers (e.g., PPARγ, adiponectin, FABP4), and smaller intracellular lipid droplets. Transmission electron microscopy (TEM) revealed IH-induced structural abnormalities, including ER fragmentation, mitochondrial disruption, nuclear enlargement, and heterochromatin formation, all of which are hallmarks of cellular aging. Furthermore, IH upregulated HIF1α, H2AX, and aging-associated histone methylation markers (H3K9me3, H3K79me3, H4K20me3), as well as extracellular matrix remodeling proteins such as fibronectin and LOX. Insulin signaling was also impaired, evidenced by decreased phosphorylation of PI3K and AKT.

Collectively, these results establish a reliable platform for long-term human adipocyte organoid culture and demonstrate its utility in modeling IH-induced adipocyte dysfunction and aging. This system offers a physiologically relevant tool for mechanistic studies and preclinical therapeutic screening targeting hypoxia-related metabolic disorders.

## Introduction

Adipose tissue is a central regulator of energy balance, metabolic homeostasis, and endocrine function ^1–5^. With advancing age and exposure to metabolic stressors, adipose tissue undergoes progressive dysfunction characterized by impaired adipogenesis ^6,7^, mitochondrial decline ^7,8^, chronic inflammation ^7,9^, and altered insulin sensitivity ^7,10^. These features contribute to systemic metabolic deterioration and increased risk of cardiometabolic diseases ^7,11,12^. While cellular and molecular mechanisms underlying adipose aging are under active investigation, suitable human models to study these processes under physiologically relevant stress conditions remain limited ^13–16^. Indeed, earlier this year the FDA and NIH each announced prioritization of new investigative and drug discovery approaches such as organoid systems and organs-on-a-chip ^17,18^.

Obstructive sleep apnea (OSA) is a prevalent and underdiagnosed condition characterized by recurrent episodes of airway collapse and reoxygenation during sleep, resulting in intermittent hypoxia (IH) ^19–24^. Growing evidence suggests that IH plays a critical role in adipose tissue dysfunction and metabolic aging ^24–26^. In both preclinical models and human studies, IH induces oxidative stress, endoplasmic reticulum (ER) stress, and pro-inflammatory signaling, all of which contribute to insulin resistance and senescence-like phenotypes in adipocytes ^24,27,28^. However, mechanistic insights into IH-induced adipocyte aging have been constrained by the absence of long-term, human-derived *in vitro* models that capture the complex interplay between hypoxia, extracellular matrix (ECM), and adipose cellular architecture.

Three-dimensional (3D) organoid technologies offer a promising platform to model human tissue biology *in vitro* ^29–33^. Organoids retain key structural and functional properties of native tissues, including cellular heterogeneity and microenvironmental interactions, and can be maintained over extended periods ^34–36^. Yet, long-term adipocyte organoids have not been widely adopted, and their utility in modeling IH-induced metabolic aging remains unexplored.

Here, we report the establishment of a robust and scalable long-term human adipocyte organoid culture system derived from stromal vascular fraction (SVF) cells ^37^ embedded in Matrigel. Organoids self-organize and mature over 10–12 days and remain viable for up to 90 days or more in culture. Using a programmable hypoxia chamber ^23,24^ to simulate clinically relevant IH conditions, we demonstrate that IH exposure suppresses adipogenesis, disrupts mitochondrial and ER integrity, promotes nuclear and epigenetic features of aging, and impairs insulin signaling. Our study provides a physiologically relevant and experimentally viable platform for investigating the mechanisms of adipocyte aging under intermittent hypoxia, with broad applications in metabolic disease research and therapeutic development.

## Materials and Methods

### Chemicals and Reagents

Unless otherwise specified, all reagents were purchased from Sigma-Aldrich, Thermo Fisher Scientific, Cayman, Inc or Cell Signaling Technology. Matrigel (Corning, growth factor-reduced), gelatin (bovine skin), and low-melting-point (LMP) agarose were used for ECM testing. Preadipocyte growth medium was obtained from ZenBio. Antibodies used for immunoblotting including anti-PPARγ, adiponectin, FABP4, Perilipin-1, FAS, ACC, HIF1α, H2AX, H3K9me3, H3K79me3, H4K20me3, fibronectin, LOX, COL11A1, VCAM-1, phospho-AKT (Ser473), phospho-PI3K, β-actin, and GAPDH (loading control) are from Cell Signaling, Sant Cruz Biotechnology, Inc and Abcam. Secondary HRP-conjugated antibodies were from Cell Signaling Technologies.

### Ethical Approval and Informed Consent

All methods were carried out in accordance with relevant guidelines and regulations. All experimental protocols were approved by the Mayo Clinic Institutional Review Board. Human adipose tissue samples from normal control subjects were collected under this approved protocol. Written informed consent was obtained from all participants prior to sample collection, in compliance with institutional ethical standards.

### Isolation of Human Subcutaneous Abdominal Fat by Biopsies

Subcutaneous abdominal fat samples were obtained from healthy adult donors (ages 18–75) under an IRB-approved protocol at Mayo Clinic per previous publications^38,39^ with minor modification. In brief, after preparation with chlorhexidine, the site was prepped and draped in a sterile manner. Local anesthesia (20cc of 1% lidocaine made to 50% solution with lactated Ringer’s) was administered subcutaneously to the area between the anterior superior iliac spine and the umbilicus. A 5-7mm incision was made before suction of subcutaneous fat was done using a 20cc syringe and fat biopsy needle, until a 1-2g sample was extracted. Local pressure was held for 5 minutes on each site. Each site was closed with 1/2“ Steri-strips and a Band-aid after povidone-iodine solution application. A wrapped bag of ice was placed on the site for one hour. Verbal and written biopsy site care instructions were given to subjects. Samples were transported in DPBS and processed within 1 hour of collection.

### Isolation of Stromal Vascular Fraction (SVF) and 2D Cell Culture

We refer to a prior published protocol by Villanueva-Carmona et al ^37^ with a slight modification. In brief, human subcutaneous fat tissues were minced and digested in collagenase type I (5mg/mL in DMEM with 5% FBS) for 45 minutes at 37°C with gentle agitation (Micro Hybridization Incubator, Robbins Scientific). The digest was filtered (100 µm strainer), centrifuged (500 ×g, 10 min), and the SVF pellet was resuspended in 5 ml ZenBio preadipocyte medium to wash one time. Cells were seeded onto collagen-coated 10cm plates and incubated for 2 days, followed by the refresh of the medium every day for the consecutive 4∼5 days to expand cells under normoxia (21% O₂, 5% CO₂, 37°C).

### Isolated SVFs Culture and Maintenance

Isolated human stromal vascular fractions (SVFs) were cultured in Subcutaneous Preadipocyte Medium (ZenBio, Cat# PM-1), supplemented with 10% fetal bovine serum (FBS) and antibiotics to support cell growth and prevent contamination. This specialized medium promotes the selective expansion of preadipocytes while maintaining their adipogenic potential. Cells were maintained under standard culture conditions (37°C, 5% CO₂) and monitored regularly for confluency and morphology. To ensure consistency and avoid phenotypic drift, only preadipocytes within passage 5 were used for downstream applications. Cells beyond passage 5 were not utilized, as extended passaging may compromise their differentiation capacity.

### 3D Organoid Formation in Matrigel and ECM Gels

SVF cells were counted with a hemocytometer cell counting reservoir under microscopy and mixed 1:1 with cold Matrigel and seeded into Biofloat U-bottom 96-well plates (5–20 µL per well, 1,250–5,000 cells/well). For ECM comparisons, gelatin (0.5%) or low-melting-point agarose gel (0.5%) were autoclaved and substituted for Matrigel. After spotting and drying the mixtures for 30 min at RT, subcutaneous preadipocyte medium was added into each well and kept at 37°C CO_2_ incubator. Organoid formation was monitored over 1–12 days using bright-field microscopy. Organoids were maintained for up to 90 days or more with medium changes every 2–3 days.

### Transmission Electron Microscopy (TEM)

Transmission Electron Microscopy (TEM) of Subcutaneous Abdominal Adipose Tissue Subcutaneous abdominal adipose tissue was embedded in agar and fixed overnight at 4°C in Trump’s fixative (1% glutaraldehyde, 4% formaldehyde in 0.1 M phosphate buffer, pH 7.2). Samples were processed using a Leica EM TP microwave processor, rinsed in phosphate buffer, and post-fixed in 1% osmium tetroxide. After washes with distilled water, samples were stained with 2% uranyl acetate, dehydrated through graded ethanol and acetone series, and infiltrated with epoxy resin before embedding in Spurr resin blocks. Ultra-thin sections (100 nm) were cut, mounted on 200-mesh copper grids, post-stained with lead citrate, and imaged using a JEM-1400 Plus TEM at 80 kV^40^.

### Adipogenic Differentiation in 2D and 3D Culture

For 2D differentiation, SVF cells were seeded in 12-well plates and induced with a standard adipogenic cocktail ^41,42^ with minor modification (containing 5 µg/ml insulin, 1µM Dexamethasone, 100µM Indomethacin, 0.5mM IBMX (3-Isobutyl-1-methylxanthine) and 10µM Rosiglitazone) for 15 days. For 3D differentiation, mature organoids (day 10–12) were exposed to the same cocktail for 3 weeks. Lipid accumulation was assessed by Oil Red O staining and quantified by dye extraction. Adipogenic marker gene expression was analyzed by Western blotting.

### Intermittent Hypoxia Exposure

Organoids and 2D cultures were placed in a programmable hypoxia chamber (BioSpherix) and exposed to IH cycles (0.1% O₂ for 30 min, followed by 21% O₂ for 30 min, cycles of 9 per day ^23,24^) continuously for up to 10–21 days or otherwise specified, simulating OSA-like IH conditions. Normoxic controls were maintained in 21% O₂.

### Western Blotting (WB)

We perform WB per our previous publications ^43–45^. In brief, cells or organoids were lysed in a lysis buffer containing 4.2M Urea, 2%SDS and 50mM Tris.HCl (pH6.8) with protease inhibitors (1 tablet from Roche in 50mL lysis buffer). Protein concentration was determined by BCA assay, and equal amounts of total proteins were loaded and separated by SDS-PAGE and transferred to PVDF membranes (Amershan^TM^Hybond^TM^, P 0.2µm; cat#10600021). Membranes were blocked in 3%∼5% nonfat milk for 30min, incubated with primary antibodies overnight at 4°C, and the probed with HRP-conjugated secondary antibodies for 1.5 hr at RT. Bands were visualized by chemiluminescence Li-COR acquisition system and quantified using Image Studio (IS). Uncropped WBs are provided in Supplementary Fig. 10.

## Statistical Analysis

All statistical analyses were performed using GraphPad Prism v. 10.0 (GraphPad Software, San Diego, CA) as per previous reports ^43,46^. Data are presented as mean ± standard error of the mean (SEM) unless otherwise stated. Comparisons between two groups were conducted using the unpaired two-tailed Student’s *t*-test. A P-value less than 0.05 was considered statistically significant.

## Results

### Development of a 3D Adipocyte Organoid Platform to Model Adipogenesis and Hypoxia-Induced Dysfunction

To model human adipose tissue *in vitro*, we isolated stromal vascular fraction (SVF) cells ^37^ from subcutaneous abdominal fat biopsies. Human SVFs contain multiple cell types, including adipose-derived stem/progenitor cells (ASCs/MSCs), endothelial lineages (both endothelial progenitors and mature endothelial cells), pericytes/mural cells, hematopoietic and immune cells (monocytes/macrophages; lymphocytes and other leukocytes), and other connective/stromal cells (fibroblasts, smooth muscle cells, preadipocytes or supra-adventitial progenitor cells) ^47,48^. The heterogeneity of SVFs makes them an ideal source for generating organoids. After five days of culture, stem cell–like clumps became visible even in 2D culture (Fig. 1A–C; Supplementary Fig.1A-D). These clumps spontaneously organized into organoid-like structures and matured into defined organoid-like clusters by days 15–20 (Fig. 1D; Supplementary Fig. 1E-H). To establish 3D adipocyte culture, SVFs were mixed with Matrigel at a 1:1 ratio, and 40–50 μL of the mixture was spotted in 12-well plates. After three weeks, well-formed adipocyte organoids were observed (Fig. 1E, F; Supplementary Fig. 2).

**Fig. 1.**
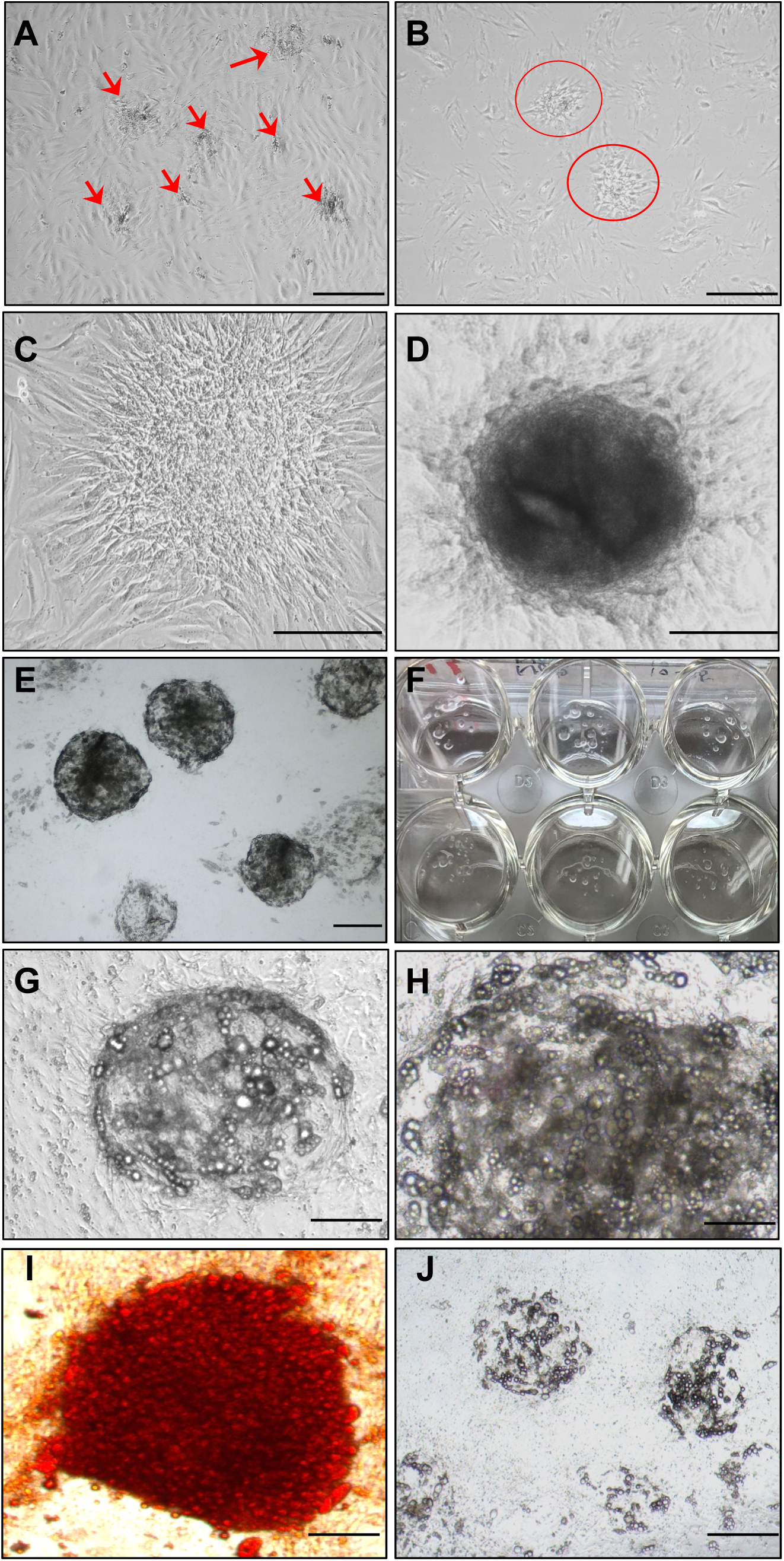
Establishment of human adipocyte organoid culture in Matrigel. (A–C) Stromal vascular fraction (SVF) was isolated from human subcutaneous abdominal fat biopsies and cultured in preadipocyte medium (ZenBio) for 5 days. Arrows in (A) indicate aggregates of stem-like cells; (B, C) show the similar aggregates at increasing magnifications. (D) Organoid-like clusters spontaneously formed in 2D culture without Matrigel. (E, F) Adipocyte organoids formed after embedding SVF in Matrigel and culturing in 12-well plates for 2 weeks; (F) shows visible organoid structures resembling fat tissue, observable by the naked eye in a 12-well plate. (G, H) Adipogenesis induced by an adipogenic cocktail for up to 3 weeks, showing progressive lipid accumulation. (I) Mature organoids stained with Oil Red O (ORO), indicating lipid-rich adipocytes. (J) Adipogenesis was markedly suppressed under intermittent hypoxia for 10 days. Scale bars: 100 µm (A, B); 150 µm (C–E, J); 200 µm (G, I); 400 µm (H).

Adipogenic differentiation using a standard induction cocktail ^41,42^ led to progressive lipid accumulation over three weeks (Fig. 1G–I), as confirmed by Oil Red O staining ^43,44^ (Fig. 1I). In contrast, organoids exposed to intermittent hypoxia (IH) for 10 days showed markedly impaired adipogenesis (Fig. 1J), indicating that adipocyte organoids are sensitive to hypoxic stress.

### Assessment of 96-Well Plates Compatibility for Adipocyte Organoid Culture

Commercially available 96-well plates, widely used in organoid culture systems such as intestinal organoids, offer standardized formats for scalable and reproducible assays ^49,50^. To adapt this approach for adipose organoid culture, we tested two commercial 96-well platforms: Biofloat U-bottom plates and Akura™ flat-bottom plates. SVF cells embedded in Matrigel successfully formed stable organoids in both systems (Fig. 2A, B).

**Fig. 2.**
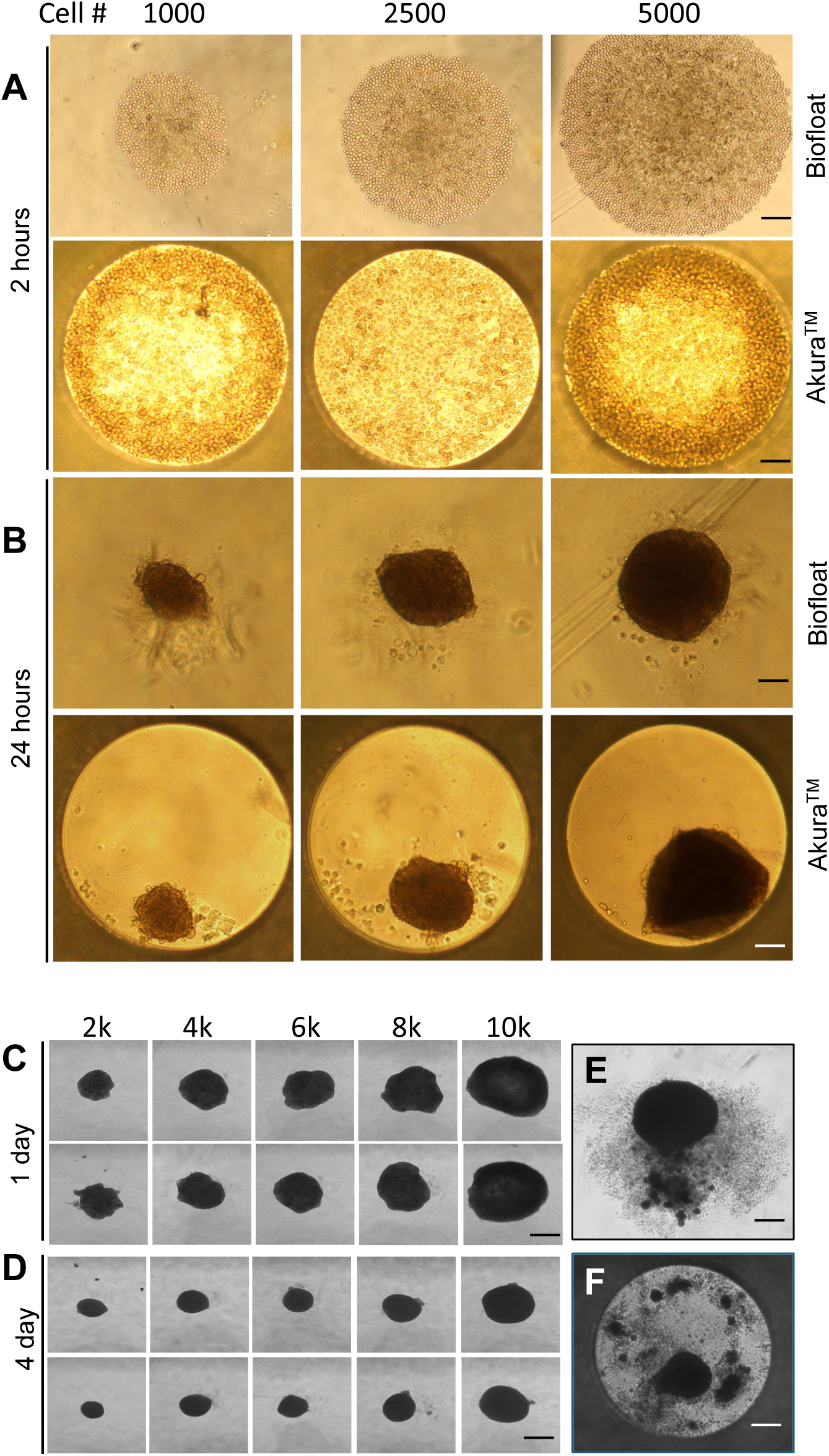
Short-term human adipocyte organoid culture from SVF in commercially available 96-well plates. SVF cells were mixed 1:1 with Matrigel and seeded at varying densities (1k, 2.5k, and 5k cells per well) into two types of 96-well plates: Biofloat U-bottom plates (Sarstedt) and Akura™ 96-well flat plates (InSphero Inc.). (A) Two hours after seeding, cells in the Biofloat plates rapidly condensed into aggregates. (B) At 24 hours post-seeding, organoid structures formed in both plate types. (C, D) Over 4 days, organoids progressively shrank across all cell densities, indicating potential alterations in cellular physiology and metabolism. (E, F) By 3 weeks, both culture systems exhibited secretion of unknown substances from the organoids, suggesting changes in their microenvironment and cell dying. Scale bars: 100 µm (A, B); 200 µm (C, D); 150 µm (E, F).

However, over time, organoids in both formats exhibited progressive shrinkage regardless of initial cell density, potentially reflecting metabolic adaptation or matrix remodeling. By three weeks, we observed secretion of unknown extracellular materials surrounding the organoids, followed by structural disassembly (Fig. 2E, F). These changes suggest a deteriorating microenvironment and compromised cell viability, possibly due to excessive mechanical forces from the hydrogel, which led to cellular malfunctions in respiration and nutrient uptake in metabolism and caused cell death. These results strongly suggest that scaffold integration into the plates is required.

### Optimization of Extracellular Support Matrices for Adipocyte Organoid Culture

A key challenge in developing physiologically relevant adipocyte organoids is the selection of an appropriate extracellular matrix (ECM) scaffold that supports 3D architecture, cell viability, and adipogenic differentiation ^51^. To systematically address this, we evaluated the effectiveness of several ECM candidates, including Matrigel, gelatin, and low-melting-point (LMP) agarose across different well plate formats to identify the most supportive microenvironment for organoid formation.

Among the tested scaffolds, Matrigel ^52^, a basement membrane extract rich in laminin, collagen IV, and entactin, proved to be indispensable (Fig. 3). When SVF cells were embedded in Matrigel and cultured in Biofloat U-bottom 96-well plates in subcutaneous preadipocyte medium with 10% FBS, they consistently formed compact, spherical, and well-structured organoids within 10–12 days (Fig. 3A; Supplementary Figs. 3-5). These organoids displayed defined borders and uniform morphology, resembling nascent adipose tissue architecture. In contrast, the same Matrigel-based cultures grown in Akura™ flat-bottom plates failed to yield comparable results, suggesting that plate geometry and surface properties significantly influence ECM behavior and cell aggregation dynamics (Fig. 3A; Supplementary Figs. 4-5).

**Fig. 3.**
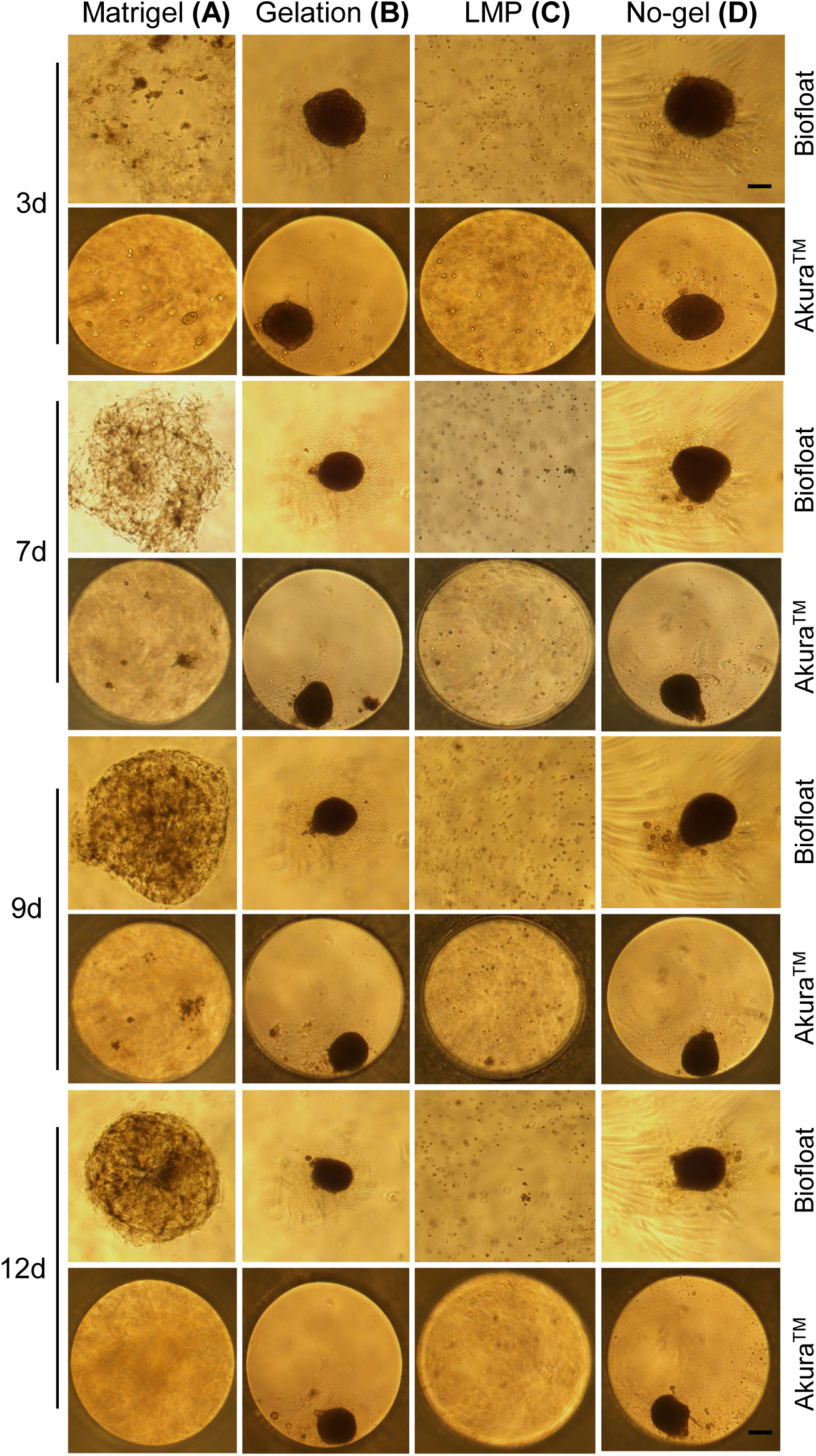
Optimization of human adipocyte organoid culture in various extracellular matrix (ECM) gels. Organoids were cultured in different ECM conditions: Gelatin (0.5%), low-melting-point agarose (LMP, 0.5%), and Matrigel using Biofloat and Akura™ 96-well plates, and monitored over a time course (days 1, 3, 5, 7, 9, and 12). Gelatin supported moderate organoid formation compared to the no-gel control, while LMP failed to support organoid formation. Matrigel consistently produced well-defined, structurally intact, and visibly mature organoids, especially by days 10–12, highlighting its superiority as a 3D matrix for adipocyte organoid culture. Scale bars represent 100 µm in all panels.

In comparison, gelatin, a collagen-derived ECM substitute, induced excessive and disorganized cell aggregation (Fig. 3B). Organoids formed in gelatin showed highly condensed clumps, even more compact than those observed in the no-gel control (Fig. 3D). This likely reflects suboptimal matrix stiffness or lack of bioactive signaling cues required for proper adipogenic morphogenesis. The resulting aggregates failed to recapitulate the natural 3D tissue architecture, highlighting gelatin’s limitations in this context.

Low-melting-point (LMP) agarose, a polysaccharide-based hydrogel commonly used for mechanical support, did not support organoid formation at all (Fig. 3C). Cells remained dispersed or formed loose without signs of organized tissue structure. LMP’s lack of cell-adhesive motifs and poor permeability for growth factors likely contributed to its failure to support adipose tissue organization.

As a comparison, we also tested advanced DMEM/F12 medium, which failed to support proper organoid formation, with poorly defined structures observed even after one month of culture compared to subcutaneous preadipocyte medium (Supplementary Fig. 6A). In addition, culturing organoids in preadipocyte medium with 0.5% FBS resulted in marked deterioration of organoid integrity compared to those maintained in 10% FBS (Supplementary Fig. 6B).

Taken together, these results underscore the critical role of Matrigel as a biologically active ECM scaffold necessary for efficient and reproducible adipocyte organoid formation. Its complex composition mimics the native extracellular environment of adipose tissue and provides the necessary biochemical and mechanical cues for cell-cell and cell-matrix interactions. Furthermore, the compatibility of Matrigel with the U-bottom well format suggests that both matrix composition and culture geometry are essential determinants of successful 3D adipose tissue modeling.

This finding establishes a foundational step for optimizing the adipocyte organoid system and sets the stage for further engineering of matrix components to enhance physiological relevance and scalability.

### Establishment of a Robust and Long-Term Adipocyte Organoid Culture System

To enable extended *in vitro* modeling of adipose tissue physiology and chronic metabolic stress, we systematically optimized a long-term adipocyte organoid culture protocol. Several key variables were tested to promote sustained viability, structural integrity, and adipogenic potential over time. Specifically, we varied Matrigel spot volumes (5–20 µL) and stromal vascular fraction (SVF) cell input densities (1,250–5,000 cells per spot) to determine the optimal conditions for organoid formation and maintenance.

Under optimized conditions, typically using 10–15 µL of Matrigel mixed with 2,500–3,500 SVF cells, compact, well-defined adipocyte organoids reliably formed within 10–12 days (Fig. 4A; Supplementary Fig. 7). These organoids displayed uniform morphology and consistent size, with well-organized cellular architecture, as confirmed by phase-contrast microscopy and histological analysis. Importantly, the optimized culture system supported stable organoid growth and viability for at least 90 days, as assessed by repeated live/dead cell staining and structural evaluation (Fig. 4B).

**Fig. 4.**
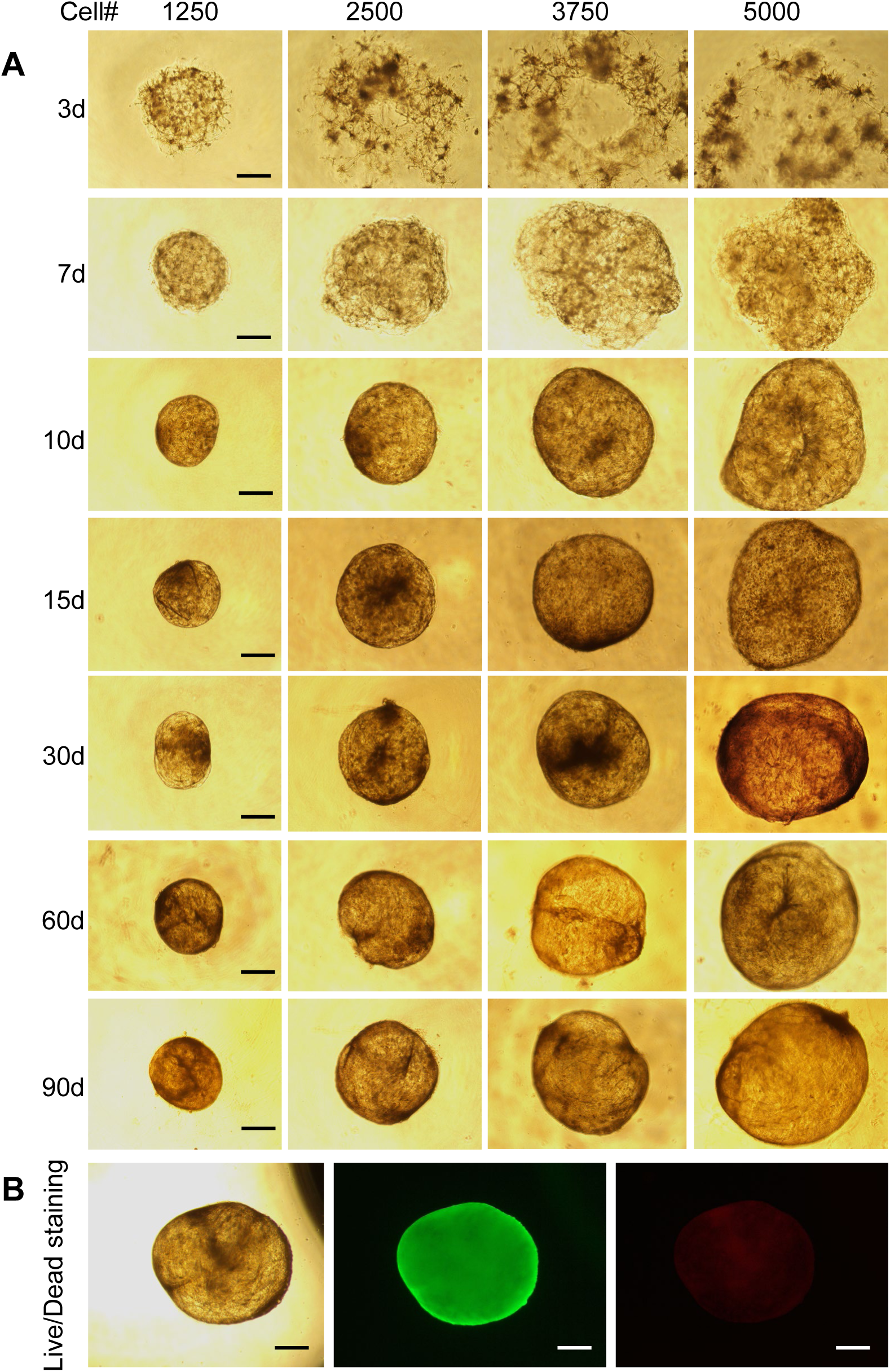
Establishment of long-term human adipocyte organoid cultures. SVF cells were mixed 1:1 with Matrigel and seeded in varying volumes (5, 10, 15, and 20 µL), corresponding to cell numbers of 1,250; 2,500; 3,750; and 5,000 per well. Organoids were cultured for up to 90 days, and an optimized protocol for long-term adipocyte organoid culture was established. (A) Organoid morphology and size became stable and consistent between days 10 and 12, and remained so through day 90. (B) Live/dead staining at day 90 demonstrated high viability, with abundant live cells (green) and only a few dead cells (red), using the Thermo Fisher Live/Dead Cell Viability Assay kit. Scale bars represent 100 µm in A and 150 µm in B.

Throughout the extended culture period, organoids retained characteristic adipocyte morphology and lipid storage capability, demonstrating the feasibility of modeling long-term adipose tissue responses *in vitro*. Unlike conventional 2D adipocyte cultures, which typically lose viability or dedifferentiate in 2-3 weeks, our 3D system preserved cellular differentiation status and tissue-like organization for several weeks to months. This advancement represents a significant step forward in adipose tissue modeling, providing a robust platform to investigate chronic metabolic insults, such as intermittent hypoxia, nutrient stress, or inflammatory stimuli, and their cumulative effects on adipocyte-associated diseases like obesity, diabetes and other cardiometabolic diseases.

Overall, this long-term organoid culture model enables mechanistic studies of adipose tissue dysfunction over extended timeframes and may serve as a valuable tool for preclinical drug testing platforms, metabolic disease modeling, and aging research.

### Intermittent Hypoxia Inhibits Adipogenesis in Organoids

Exposure to intermittent hypoxia (IH) for 2–3 weeks markedly impaired adipogenic differentiation in organoids (Fig. 5). Morphologically, IH-treated organoids secreted unknown extracellular substances surrounding organoids (Fig. 5A, B) and exhibited a notable reduction in lipid accumulation (Fig. 5C–F). This result is further confirmed in 2D preadipocyte culture induced by adipogenic cocktail (Supplementary Fig. 8). Western blot analysis demonstrated downregulation of key adipogenic markers, including PPARγ, adiponectin, FABP4, perilipin-1, ACC, and FAS (Fig. 5G). Transmission electron microscopy (TEM) further revealed smaller and fewer lipid droplets in IH-exposed organoids *versus* normoxia (Fig. 5H, I), consistent with compromised lipid storage and impaired adipogenesis in our initial experiments (Fig. 1G-J).

**Fig. 5.**
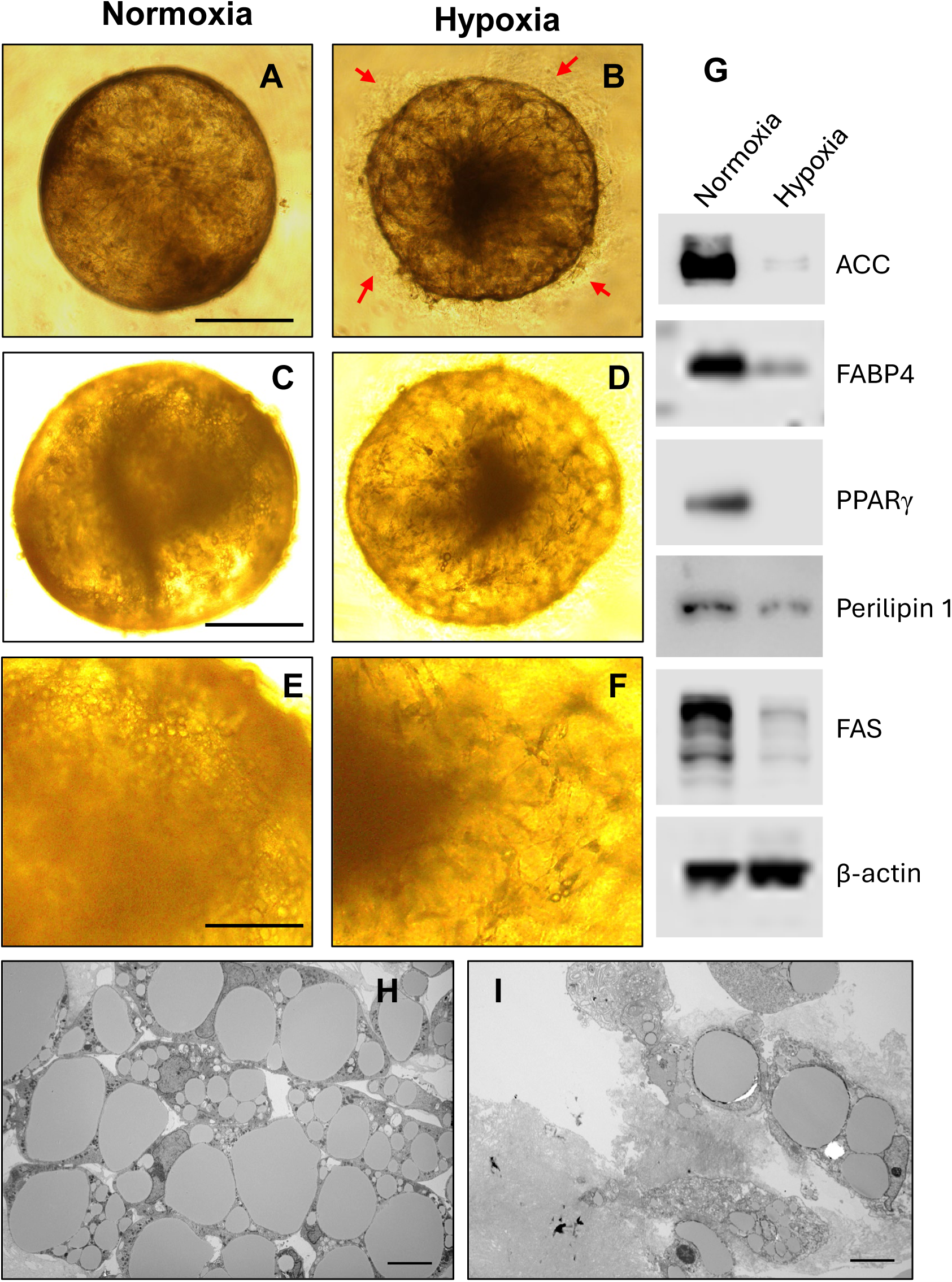
Intermittent hypoxia (IH) inhibits adipogenesis in human adipocyte organoids. (A–B) Compared to normoxic conditions (A), IH treatment for 2 weeks led to an increase in the secretion of unknown extracellular substances surrounding the organoids (B). (C–F) IH significantly impaired adipogenesis during a 3-week induction period using an adipogenic cocktail. E&F is the amplified magnification for C&D, respectively. (G) Gene expression analysis confirmed that IH reduced the expression of key adipogenic markers, including ACC, Adiponectin, FABP4, PPARγ, Perilipin-1, and FAS. (H, I) Transmission electron microscopy (TEM) revealed that IH reduced both the number and size of intracellular lipid droplets, further supporting impaired adipogenesis under hypoxic conditions. Scale bars: 100 µm (A, B); 150 µm (C, D); 400 µm (E, F); 1 µm (H, I).

### IH Induces Organelle Damage and Aging-Associated Structural Changes

TEM analysis provided ultrastructural insights into the impact of intermittent hypoxia (IH) on adipocyte organoids. Compared to normoxic controls, IH-exposed organoids exhibited significant disruption of intracellular architecture. Notably, the endoplasmic reticulum (ER) appeared dilated and fragmented, indicating ER stress and potential impairment in protein folding and lipid biosynthesis (Fig. 6A–D). This structural alteration suggests a maladaptive response to hypoxic conditions, which may compromise adipocyte function and viability.

**Fig. 6.**
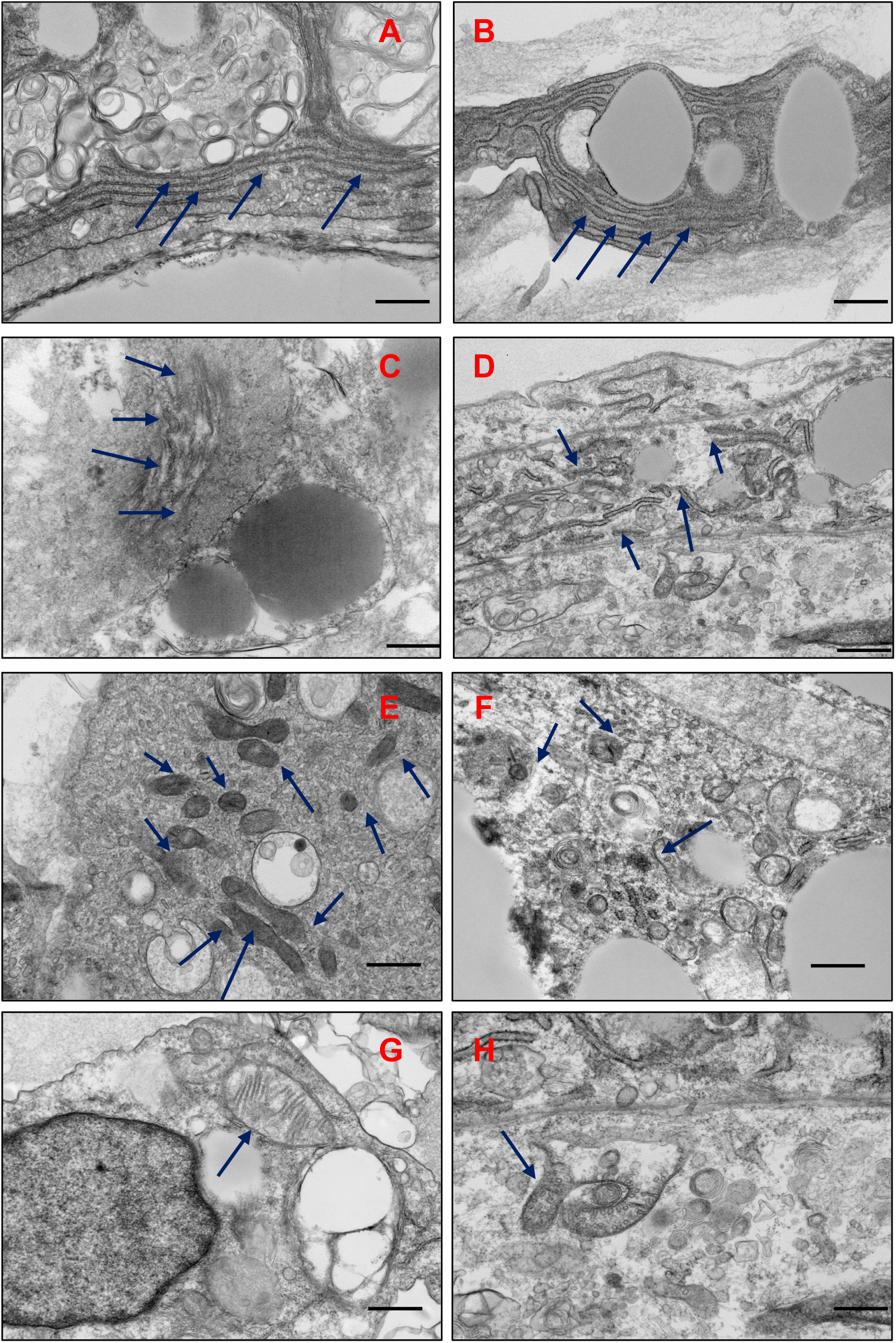
Intermittent hypoxia (IH) induces ER disruption, mitochondrial dysfunction, and aging-associated features in adipocyte organoids. (A, B) Under normoxic conditions, organoids exhibit well-organized, intact endoplasmic reticulum (ER) structures with well-formed linear morphology. (C, D) IH treatment results in dilated, fragmented, and disconnected ER, indicating ER stress and structural damage. (E, F) IH reduces mitochondrial abundance within organoid cells. (G, H) Mitochondria under IH show irregular morphology (H), including smaller size and disrupted cristae, compared to the normal elongated structure seen in normoxia (G). These structural changes suggest increased oxidative stress, mitochondrial dysfunction and may reflect early signs of cellular aging. Scale bars: 1 µm (A–F); 500 nm (G, H).

In addition to ER abnormalities, IH markedly reduced mitochondrial abundance (Fig. 6E, F). Remaining mitochondria were visibly smaller and displayed disorganized or swollen cristae, a hallmark of mitochondrial dysfunction (Fig. 6G, H). Quantitative morphometric analysis of mitochondria showed significant differences between normoxic and hypoxic fat organoids. Under normoxic conditions, mitochondria possessed a larger mean area (1404 ± 259 nm²) and perimeter (6.78 ± 0.21 nm) when compared to their counterparts under hypoxic conditions, where mitochondria recorded considerably smaller area (476 ± 118 nm²) and reduced perimeter (4.03 ± 0.96 nm). These changes are consistent with impaired oxidative metabolism and suggest an early onset of mitochondrial deterioration, which is often associated with cellular aging and metabolic stress. Furthermore, the plasma membrane appears smooth and continuous, maintaining stable cell–cell contacts with well-defined intercellular junctions in normoxic conditions, thereby preserving tissue integrity. In contrast, hypoxic conditions disrupt membrane integrity, leading to irregularities and less distinct junctions between adjacent cells (Supplementary Fig. 9). These alterations reflect hypoxia-induced membrane leakage and weakened intercellular junctions. Collectively, these findings highlight that IH not only impairs adipogenic differentiation but also induces profound subcellular remodeling, reflective of cellular stress and degeneration.

### Nuclear Alterations and Epigenetic Remodeling Under IH

Intermittent hypoxia (IH) induced striking nuclear changes in adipocyte organoids, indicative of structural stress and early senescence. TEM in high-resolution imaging revealed enlarged nuclei with irregular shapes, increased chromatin condensation along the nuclear periphery, and thinning or focal discontinuities of the nuclear envelope (Fig. 7A–D). These nuclear abnormalities closely resemble classical features of cellular aging and senescence, including altered chromatin architecture and compromised nuclear integrity.

**Fig. 7.**
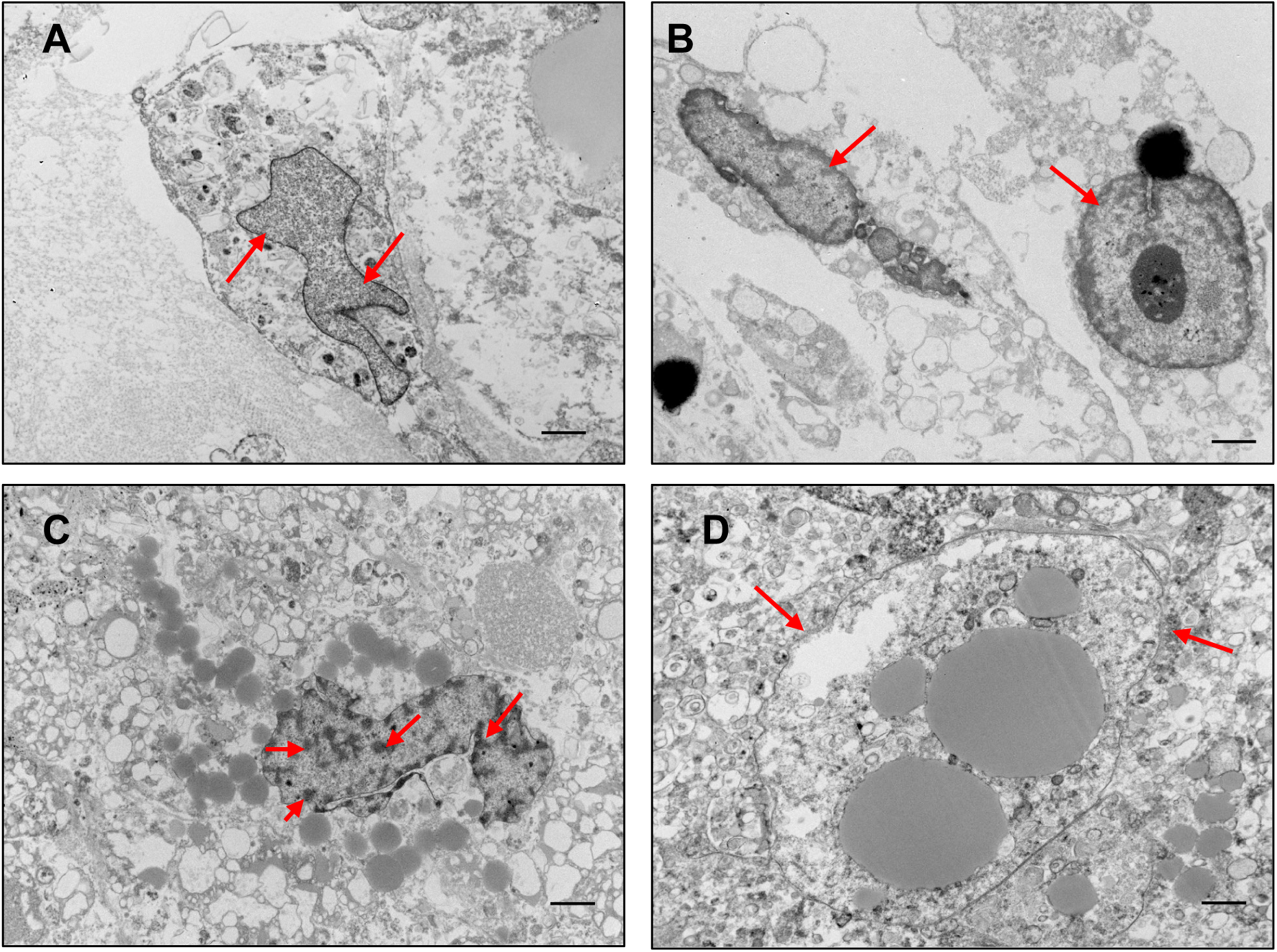
Intermittent hypoxia (IH) induces nuclear enlargement, inflammation, and heterochromatin formation—hallmarks of cellular aging. (A) Under normoxic conditions, organoid nuclei exhibit normal morphology with intact nuclear membranes and evenly distributed chromatin, as observed by TEM. (B–D) IH exposure leads to pronounced nuclear enlargement and signs of inflammation (B), with chromatin condensation along the nuclear periphery forming distinct heterochromatin (C), and visible thinning or damage to the nuclear membrane (D). These nuclear alterations are consistent with aging-associated cellular stress and epigenetic reprogramming. Scale bars represent 2 µm in all panels.

To further investigate the molecular mechanism of these morphological changes, we examined key markers of hypoxia, DNA damage, and epigenetic remodeling. IH exposure led to a marked upregulation of hypoxia-inducible factor 1-alpha (HIF1α), a central regulator of the cellular response to oxygen deprivation (Fig. 8A). In parallel, there was an increase in H2AX, a well-established marker of DNA double-strand breaks, indicating elevated genotoxic stress under IH conditions (Fig. 8A, B).

**Fig. 8.**
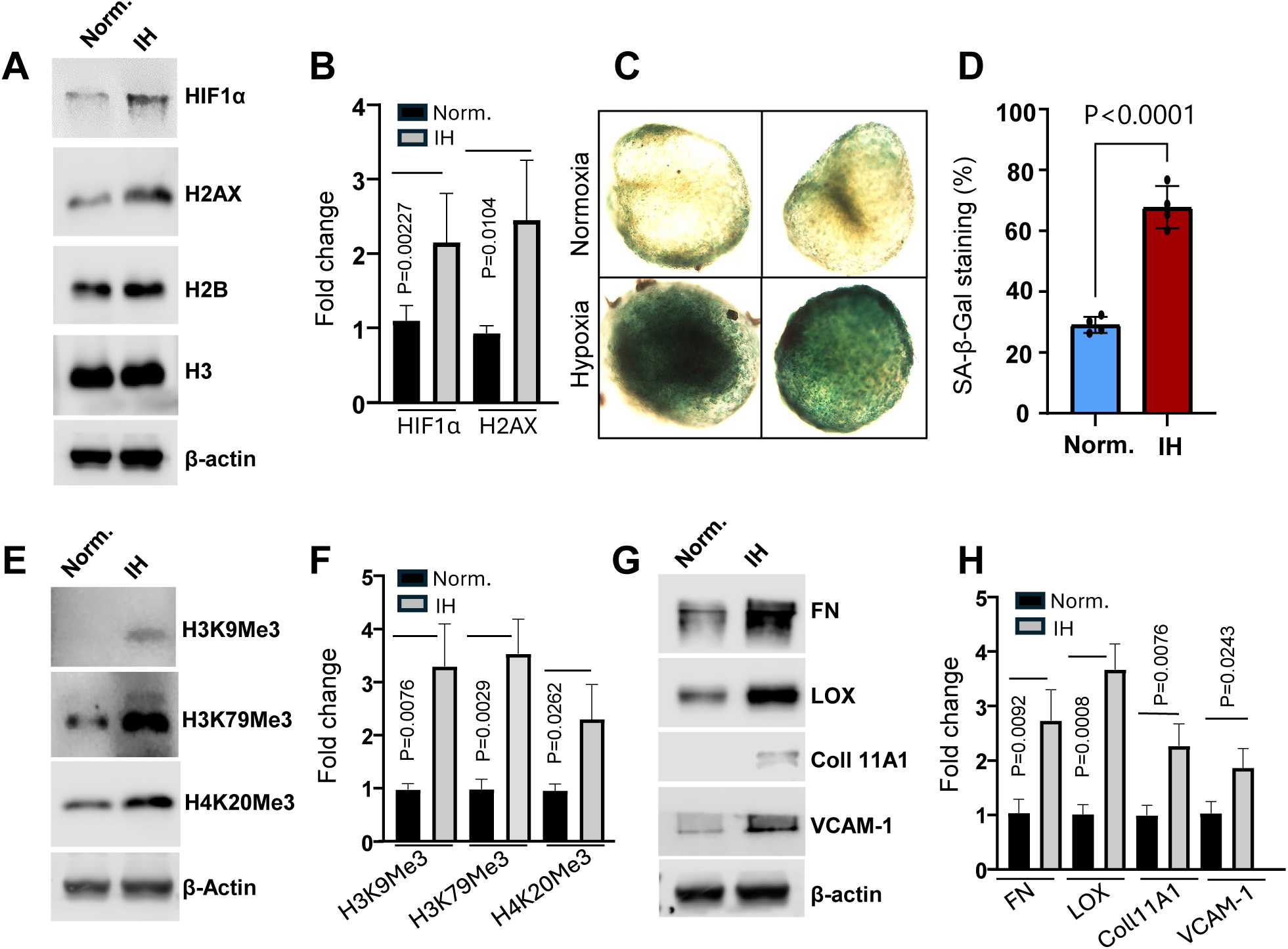
Intermittent hypoxia (IH) upregulates HIF1α, senescence markers, histone methylation, and ECM remodeling proteins. (A, B) IH treatment for 10 days significantly increased expression of HIF1α and H2AX, both established markers of cellular senescence and aging. (C, D) IH treatment for 7 days markedly increased cellular senescence, as shown by SA-β-gal staining (C). Quantification is presented in (D) (Student’s t-test, n = 4 organoids per group, P < 0.0001). (E, F) IH also elevated levels of aging-associated histone methylation marks, including H3K9me3, H3K79me3, and H4K20me3, indicating epigenetic reprogramming. (G, H) Expression of extracellular matrix (ECM) remodeling proteins: fibronectin (FN), lysyl oxidase (LOX), and collagen type XI alpha 1 (COL11A1), as well as the adhesion molecule VCAM-1 was upregulated under IH, suggesting ECM and microenvironmental alterations that may contribute to aging and inflammation.

Importantly, IH also triggered significant changes in the epigenetic landscape of the organoids. Western blot analysis and immunostaining revealed increased levels of repressive histone methylation marks, including trimethylated histone H3 lysine 9 (H3K9me3), lysine 79 (H3K79me3), and histone H4 lysine 20 (H4K20me3) (Fig. 8C–D). These histone modifications are known to be associated with heterochromatin formation, transcriptional repression, and epigenetic aging. Their accumulation under IH suggests a global chromatin reorganization toward a more repressive state, potentially contributing to reduced transcriptional plasticity, impaired adipogenic gene expression, and senescence-like phenotypes.

Together, these findings highlight that IH not only disrupts nuclear architecture but also induces profound epigenetic remodeling, recapitulating key features of premature aging in adipose tissue. This establishes a mechanistic link between hypoxic stress and the decline of adipocyte function through nuclear and chromatin-level dysregulation.

### ECM Remodeling and Insulin Resistance Induced by Intermittent Hypoxia (IH) During Adipogenesis

Further molecular and protein analyses revealed significant upregulation of extracellular matrix (ECM) remodeling components in cells exposed to IH during adipogenesis. Specifically, key ECM-associated proteins such as fibronectin, lysyl oxidase (LOX), and collagen type XI alpha 1 chain (COL11A1) were markedly elevated, along with the upregulation of adhesion molecule VCAM-1 (Fig. 8E, F). These changes suggest that IH disrupts the structural and biochemical composition of the adipogenic niche, potentially altering cell–matrix interactions and tissue stiffness. Such remodeling of the ECM may negatively impact adipocyte differentiation and function, further contributing to metabolic dysregulation. In parallel, we observed a suppression of insulin signaling pathways under IH conditions. Specifically, the phosphorylation levels of PI3K and AKT, critical mediators of insulin action, were significantly reduced upon adipogenic stimulation (Fig. 9), indicating that IH impairs the normal insulin responsiveness of differentiating adipocytes. Together, these findings highlight a dual impact of IH on both ECM remodeling and insulin signaling, providing mechanistic insight into how IH contributes to adipose tissue dysfunction, abnormal metabolism and insulin resistance.

**Fig. 9.**
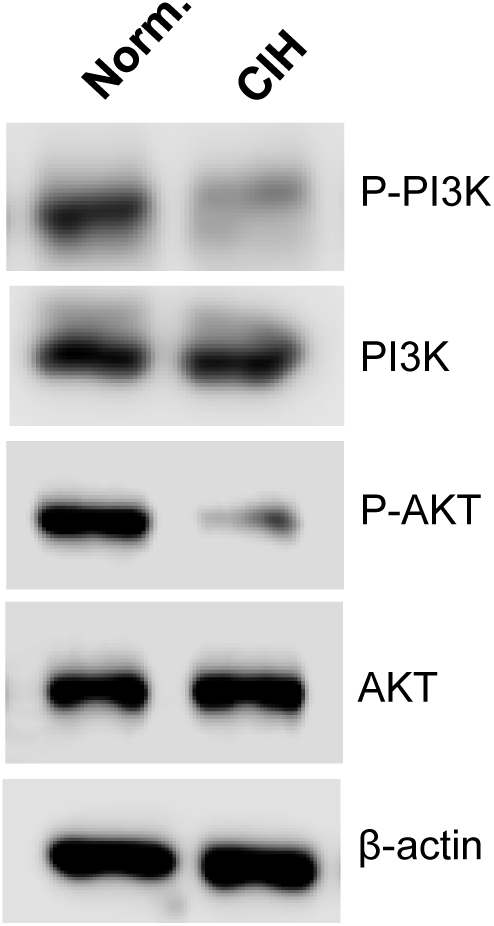
Intermittent hypoxia (IH) impairs insulin signaling during adipogenesis in human adipocyte organoids. Insulin signaling was assessed under normoxic and IH conditions during adipogenic differentiation. IH-treated organoids showed significantly reduced phosphorylation of PI3K and AKT, key mediators of insulin signaling, indicating impaired insulin responsiveness and altered metabolic function. These findings suggest that IH induces insulin resistance in adipocyte organoids. Results are representative of four independent experiments.

## Discussion

Recent advances in adipose organoid culture have significantly improved our ability to model adipose tissue physiology *in vitro*. Notable progress includes scaffold-free organoid generation using magnetic bioprinting, which enables the maintenance of late-stage adipocyte characteristics while minimizing buoyancy-related instability ^53^. High-throughput production of uniformly sized spheroids through microfluidic encapsulation has also expanded the potential for drug screening applications ^54^. Moreover, efforts to incorporate vascular components—such as the use of human microvessel fragments ^55^ or ruminant-derived stromal vascular fractions ^56^ have enhanced the physiological relevance of these models by supporting angiogenesis and adipocyte–endothelial interactions.

Some organoid systems have even captured immune-metabolic crosstalk by retaining resident immune cells ^57^, while others have succeeded in generating beige adipose organoids capable of inducible thermogenesis and endocrine signaling ^58^. However, despite these encouraging developments, most current adipocyte organoid platforms are optimized for short-to mid-term culture (typically 2–4 weeks) and do not model the chronic progression of metabolic disorders such as cardiovascular disease, insulin resistance, or lipotoxicity.

This limitation presents a major barrier to studying long-term cellular remodeling or therapeutic intervention in disease-relevant settings. Scaffold-based models still suffer from variability in extracellular matrix (ECM) composition, and reproducibility remains a concern ^59,60^. In contrast, our current system addresses several of these limitations by offering longer-term culture, maintaining functional adipocyte markers and viability, and mimicking relevant stimuli outcomes like hypoxia or nutrient excess.

Together, these findings highlight the importance of developing more physiologically and pathologically relevant adipose organoid systems that can capture the dynamic and chronic nature of adipose tissue dysfunction in obesity and related metabolic diseases.

In this study, we established and validated a robust, scalable, and long-term human adipocyte organoid culture system capable of modeling intermittent hypoxia (IH)-induced adipocyte dysfunction and aging. By embedding human stromal vascular fraction (SVF) cells in Matrigel and optimizing culture conditions in commercially available Biofloat U-bottom plates, we achieved reproducible adipocyte organoid formation within 10–12 days and, critically, maintained their viability and structural integrity for up to 90 days or more (Fig. 4). To our knowledge, this represents the longest functional culture of human adipocyte organoids reported to date. Most existing adipose organoid systems support differentiation and maintenance for only 2–3 weeks, limiting their utility for modeling chronic metabolic conditions such as cardiovascular disease (CVD), obesity, and type 2 diabetes, which develop chronically and over decades ^61–63^. The ability to sustain long-term adipocyte physiology *in vitro* fills a major technological gap and enables extended temporal analysis of molecular mechanisms and drug responses.

A second key innovation of our study is the integration of a programmable hypoxia chamber to simulate OSA-relevant IH conditions in a human fat organoid model. OSA affects nearly one billion individuals globally and is increasingly recognized as a contributor to accelerated aging and cardiometabolic dysfunction ^64^. However, the cellular mechanisms linking IH to adipocyte aging remain poorly understood, due in part to the lack of human-specific *in vitro* models. By exposing organoids to controlled IH cycles, we recapitulated hallmark features of aging, including suppressed adipogenesis, mitochondrial fragmentation, ER stress, nuclear chromatin condensation, and upregulation of senescence-associated histone methylation and ECM remodeling proteins (Fig. 8). These findings position our model at the forefront of OSA-aging research and provide a powerful experimental system to dissect the pathophysiological impact of IH on human adipose tissue.

Importantly, our fat organoid system offers strong translational value for drug discovery. Unlike immortalized cell lines or short-lived 2D primary cultures, our long-lived organoids maintain cellular viability, metabolic function, and structural fidelity for months (Fig. 4; Supplementary Fig. 7). This is particularly advantageous for evaluating chronic effects of candidate compounds targeting obesity, insulin resistance, or age-related adipose degeneration. Furthermore, the compatibility of our organoid model with 3D bioprinting platforms opens new opportunities to develop customizable and high-throughput organoid arrays for automated drug screening. The spatial control enabled by 3D bioprinting, combined with the stability of our long-term culture system, provides an ideal framework for testing drug efficacy and toxicity under physiologically relevant conditions. As pharmaceutical development increasingly requires robust human-based preclinical models, our platform offers a timely and scalable solution for bridging mechanistic discovery with therapeutic innovation.

In summary, this study introduces a novel long-term 3D adipocyte organoid platform that models human adipose aging under intermittent hypoxia. Our findings not only offer new mechanistic insights into OSA-induced adipocyte dysfunction but also establish a foundational tool for studying chronic metabolic diseases and accelerating drug development through advanced organoid technologies such as 3D bioprinting. Future applications of this system may identify new therapeutic targets and transform preclinical strategies for metabolic and aging-related disorders.

## Acknowledgements

We gratefully acknowledge our collaborators at Mayo Clinic for their valuable discussions, insights, and technical support throughout this study. We also thank the Mayo Clinic Microscopy and Imaging Core, Histology Core, and Biorepository Core for their expert assistance and contributions to data acquisition and analysis.

## Author Contributions

V.K.S. and Y.D. conceptualized and initiated the project. Y.D. performed the majority of the experiments and drafted the initial manuscript. V.K.S. and Y.D. jointly revised and finalized the manuscript. J.W., F.F. and I.H.A. contributed to selected experiments and imaging analyses. S.K. and J.B. recruited subjects and performed the subcutaneous abdominal fat biopsies.

## Sources of Funding

This research was supported by NIH grant HL65176 (VKS) and by a grant from the Corinne and William Little Foundation to the Mayo Clinic. JMB was supported by NHLBI T32-HL007111 as well as NIA K12AR084222 and U54AG044170.

## Disclosure of conflicting interests

VKS serves as a consultant for Axsome, Jazz Pharmaceuticals, Lilly, Mineralys, iRhythm, and ApniMed, and is a member of the Scientific Advisory Board for Sleep Number. The authors report no other conflicts of interest.

## Data availability

The data supporting this study will be available from the corresponding author upon publication.

## Supplementary supporting information

Additional support information can be found online at xxxxxx.

