## supplementary Figures for "Development of Long-Term Human Adipocyte Organoids Manifesting Aging in Response to Intermittent Hypoxia"

Runing title: Long-Term Human Adipocyte Organoids Mimicking Hypoxic Aging

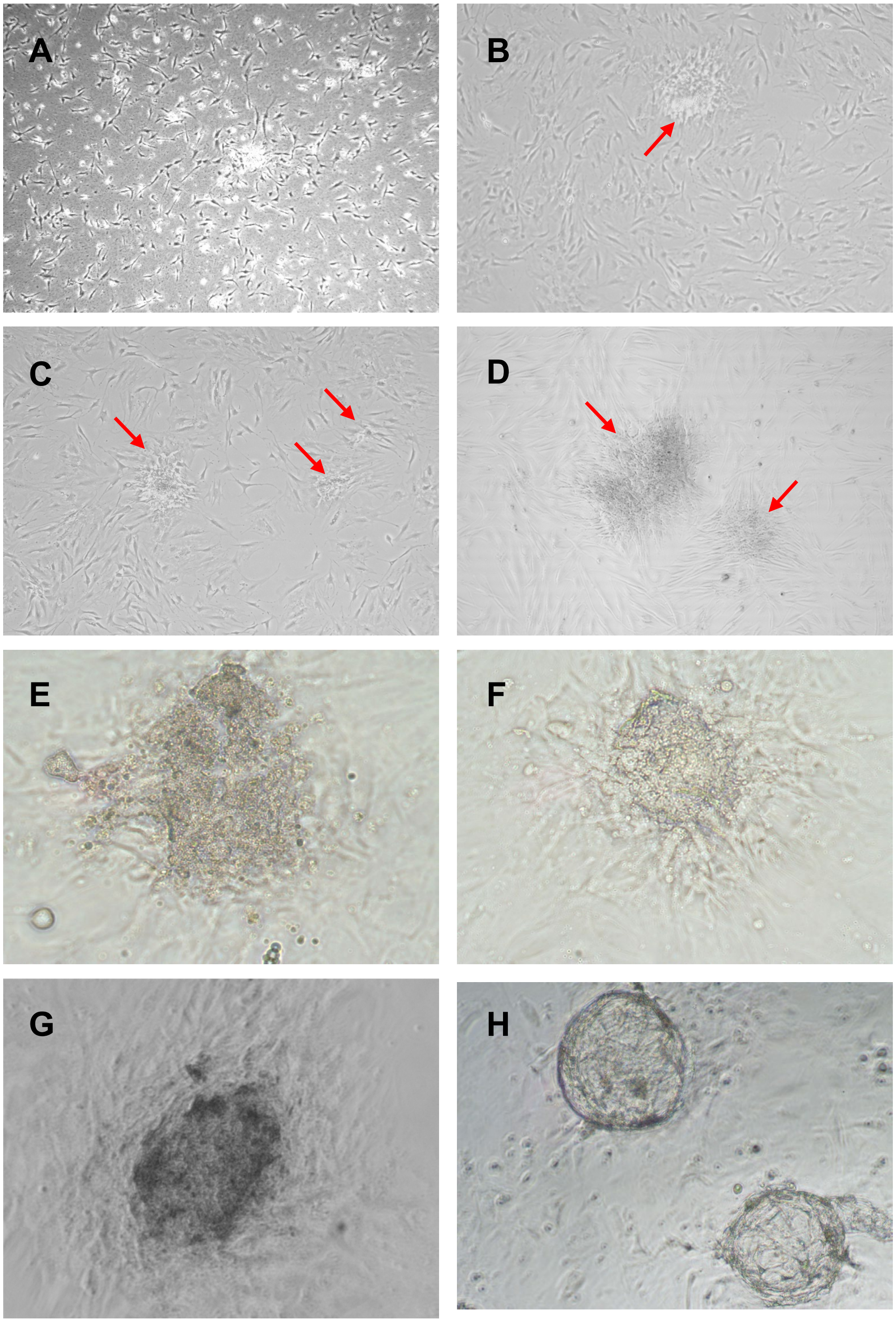


**Supplementary Fig. 1. Observation of human adipocyte organoid-like structure formation in 2D culture without Matrigel.**

SVF was isolated from human subcutaneous abdominal fat and cultured under standard 2D conditions. The culture revealed spontaneous aggregation of stem-like cells and gradual formation of fat organoid-like structures over time.

(A) Human SVF cultured for 5 days after removal of debris by triple aspiration for 3 days.

(B) Arrow indicates clusters of stem-like cells.

(C, D) Additional views showing larger clumps of stem-like cells.

(E, F) Gradual formation of organoid-like structures observed by day 15 in 2D culture.

(G, H) Organoid-like structures can form naturally even in 2D culture conditions without the use of Matrigel or 3D scaffolding.

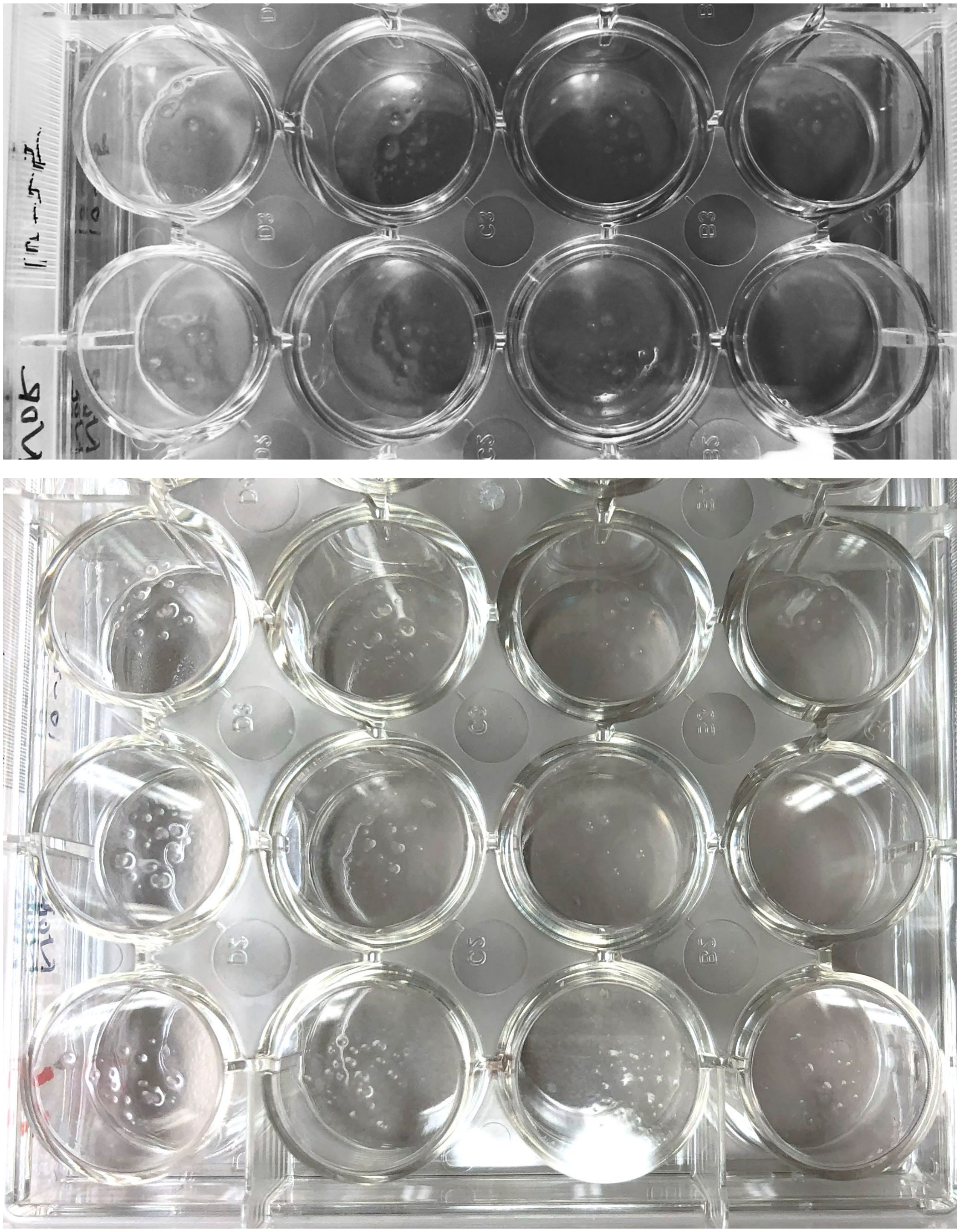

**Supplementary Fig. 2. Formation of human adipocyte organoids in 3D Matrigel cultures.**
SVF cells embedded in Matrigel and cultured in regular 12-well flat plates developed visible fat organoids over 3~4 weeks. Mature organoid structures were easily observed by the naked eye.

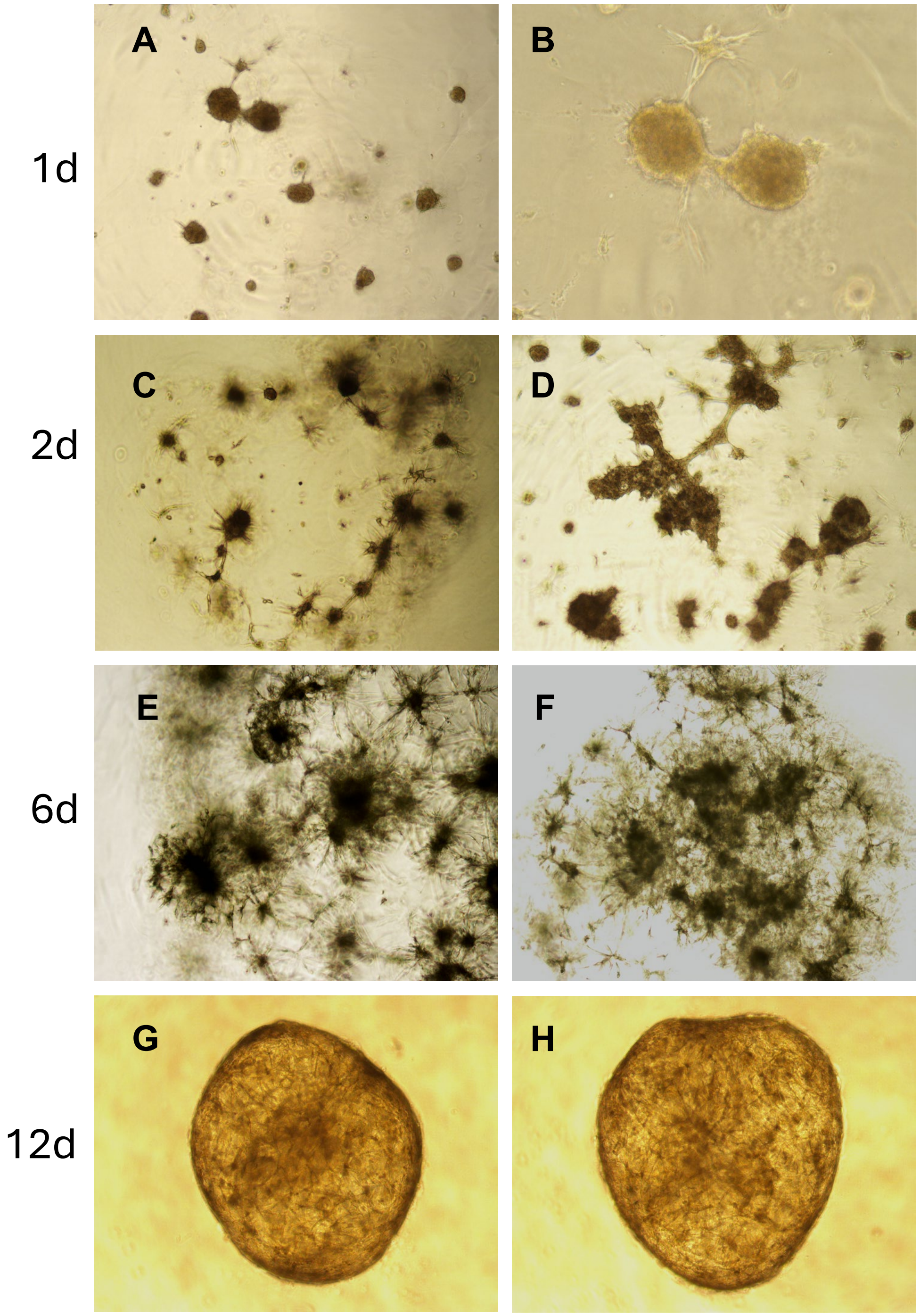


**Supplementary Fig. 3. Human adipocyte organoid formation over 10–12 days in Matrigel.**
Human SVF cells were mixed 1:1 with Matrigel and seeded into 96-well Biofloat U-bottom plates (Sarstedt, Inc). Cultures were monitored over a time course from day 1 to day 12, with each well seeded with 2,500 SVF cells. From day 1, aggregates of stem-like cells rapidly condensed into compact clusters. By days 10–12, well-structured and morphologically mature adipocyte organoids had consistently formed within the Matrigel scaffold, demonstrating the supportive role of the 3D ECM microenvironment in organoid development.

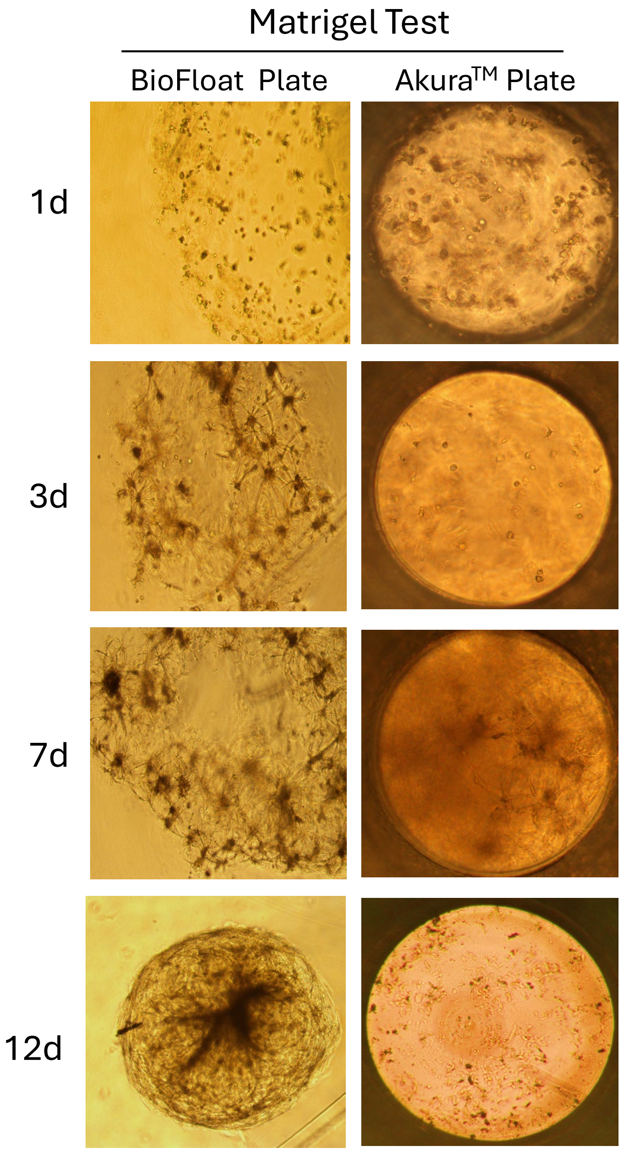


**Supplementary Fig. 4. Human adipocyte organoid formation occurs only with the combination of Biofloat plates and Matrigel, but not in Akura™ plates.**

SVF cells were mixed 1:1 with Matrigel and seeded into either Biofloat U-bottom plates or Akura™ flat-bottom 96-well plates. Organoid development was monitored over 12 days.

Results showed that organoid formation occurred consistently in Biofloat plates, with mature structures appearing by days 10–12, whereas no organoids formed in or Akura™ flat-bottom plates under the same conditions.

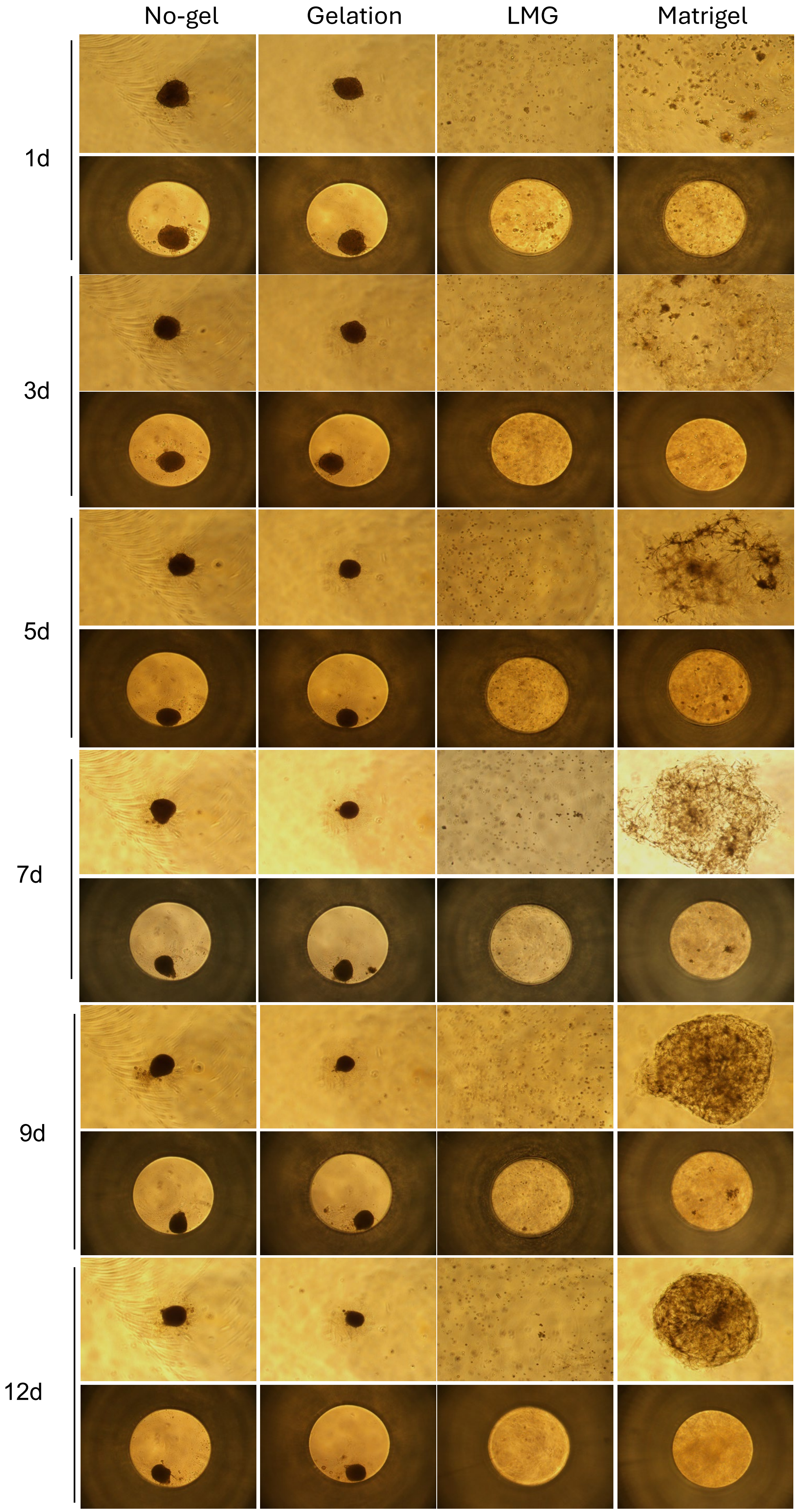


**Supplementary Fig. 5. Optimization of human organoid culture in ECM-based scaffold gels (see next page).**

**Supplementary Fig. 5. Optimization of human organoid culture in ECM-based scaffold gels.** We evaluated various extracellular matrix (ECM) substrates for supporting human organoid formation, including gelatin (0.5%), low-melting agarose gel (LMG, 0.5%), and Matrigel. All gels were mixed cells at 1:1. Organoid development was monitored over a time course (days 1, 3, 5, 7, 9, and 12). We found that only Matrigel, when combined with Biofloat U-bottom 96-well plates, consistently supported the formation of well-structured organoids within 10–12 days. In contrast, Akura™ flat-bottom 96-well plates failed to support proper organoid formation, and LMG did not support organoid formation at all. Although gelatin promoted tight cell aggregation, this excessive compaction appeared detrimental to normal cellular physiology and metabolic function. In each panel, upper panel: Biofloat U-shape 96-well plates; lower panel: Akura™ flat 96-well plates.

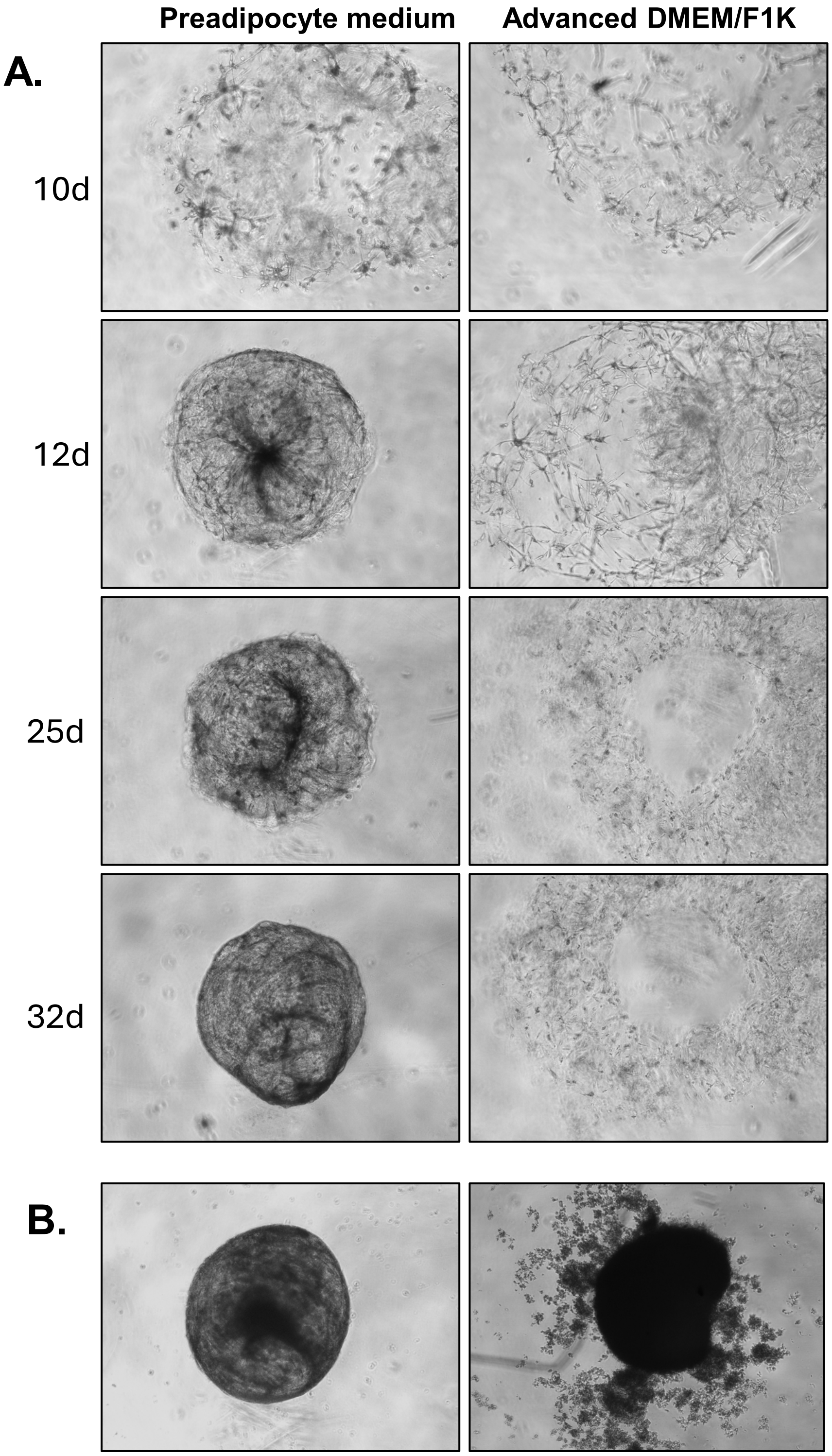


**Supplementary Fig. 6. Optimization of human adipocyte organoid culture in different media.**
Organoids were cultured in Biofloat U-bottom 96-well plates seeded with 2,500 SVF cells per well.
(A) Comparison of media showed that commercially available "Advanced" medium produced poorly structured, fragile organoids over one month, whereas the preadipocyte medium (ZenBio) specifically formulated for adipocyte cultures supported robust and healthy organoid formation within 10-12 days.
(B) To assess the requirement for serum, long-term cultures (>30 days) were established with 10% FBS and lower FBS (0.5%) supplementation. In the 0.5% FBS preadipocyte culture medium, organoids gradually deteriorated, disassembled, and exhibited loss of viability after approximately one month.

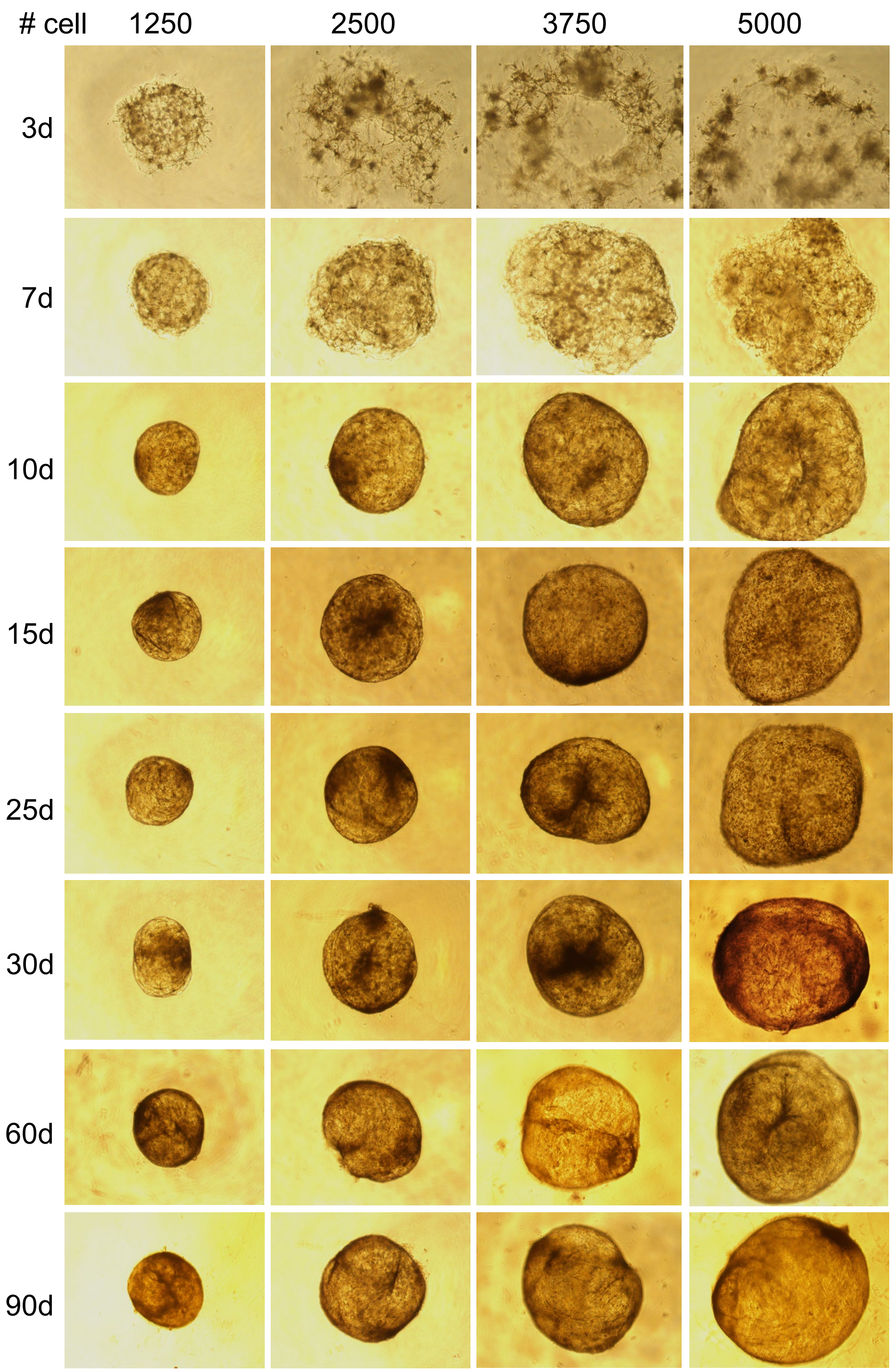


**Supplementary Fig. 7. Establishment of long-term fat organoid culture-time course analysis.** Stromal vascular fraction (SVF) cells were mixed with Matrigel at a 1:1 ratio and seeded as spot organoids in volumes of 5 µL, 10 µL, 15 µL, and 20 µL, corresponding to cell densities of 1,250; 2,500; 3,750; and 5,000 cells, respectively. Organoids successfully formed within 10–12 days and maintained stable morphology and size for up to 3 months in culture. These results demonstrate the successful establishment of a robust long-term fat organoid culture system.

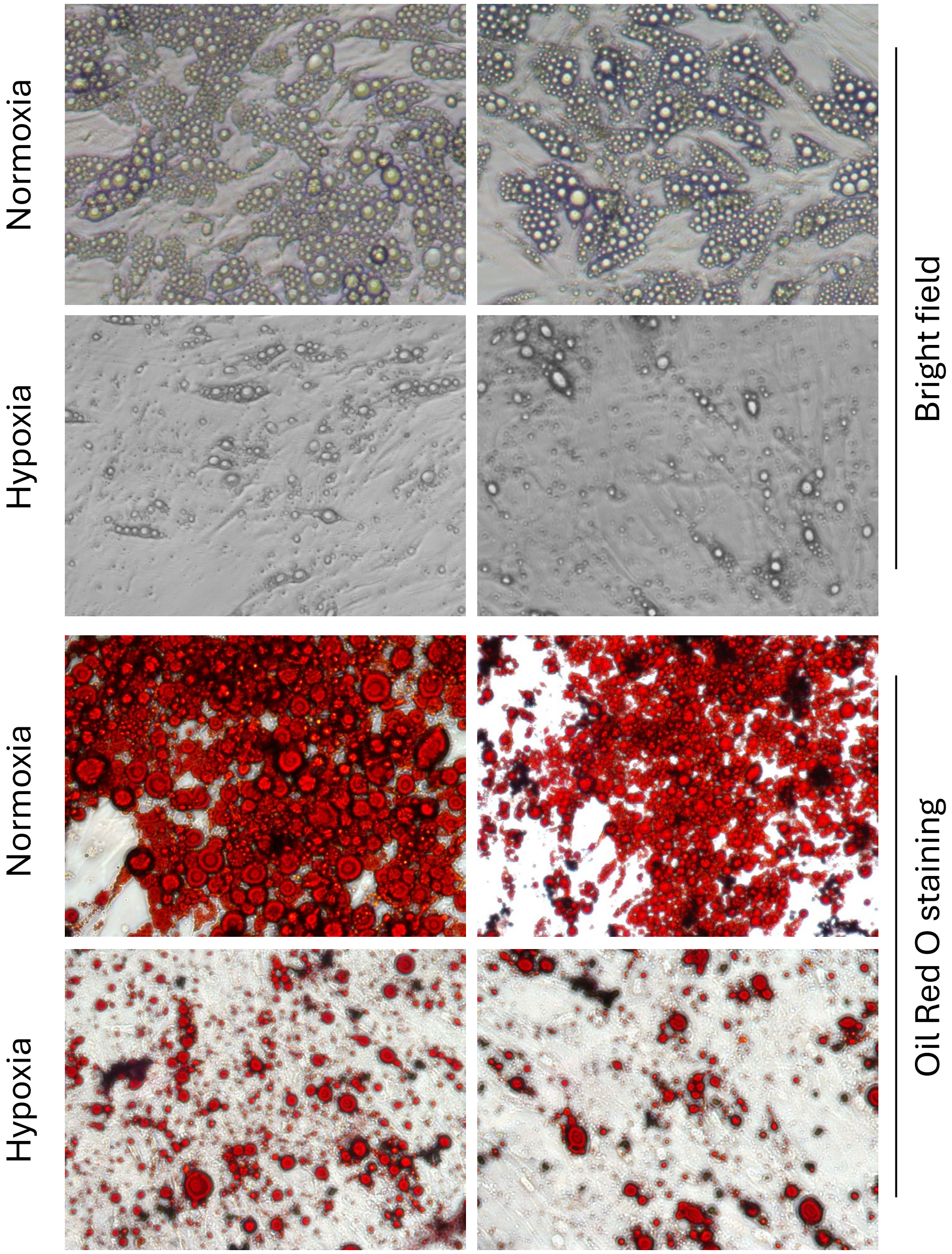


**Supplementary Fig. 8. Intermittent hypoxia (IH) impairs adipogenesis in 2D preadipocyte culture.** To complement our findings from 3D organoid models, we conducted adipogenic differentiation in a 2D culture system. Under normoxic conditions, human preadipocytes underwent robust adipogenesis after 15 days of exposure to a standard differentiation cocktail revealed by Oil Red O staining. In contrast, IH treatment markedly inhibited adipocyte differentiation, indicating a suppressive effect of IH on adipogenesis in both 2D and 3D culture systems.

### Plasma Membrane and Cell-Cell Junctions

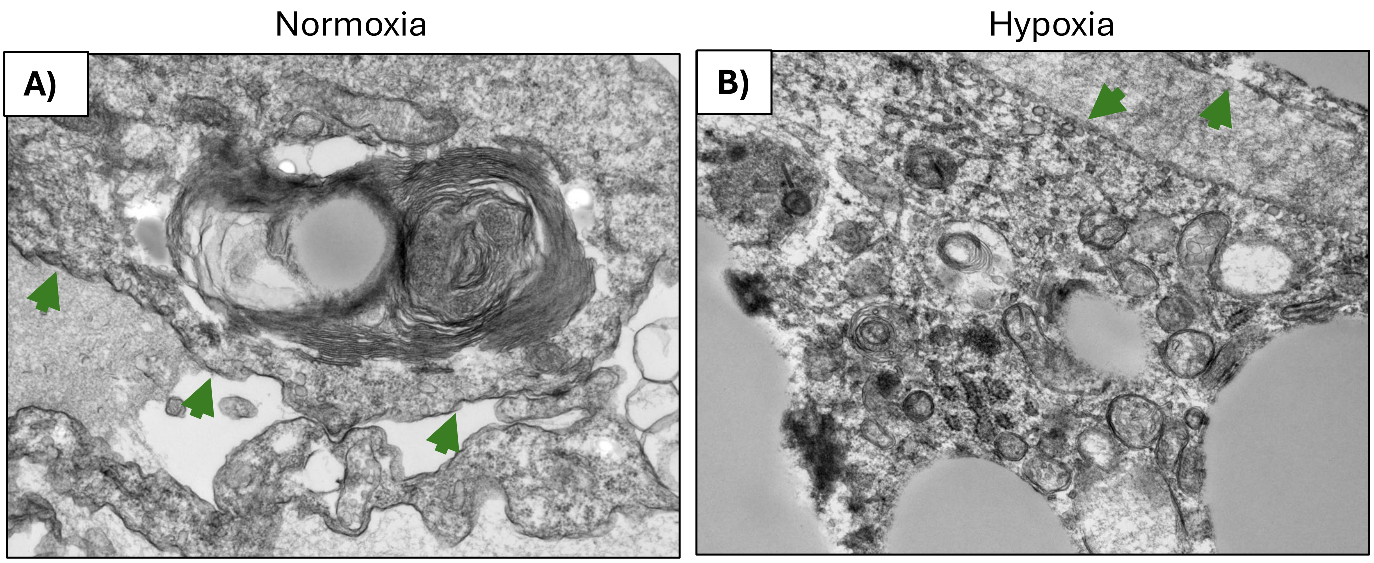


**Supplementary Fig. 9. Investigation of ultrastructural features of plasma membrane and cell-cell junctions under both normoxic and hypoxic conditions.**
Representative TEM images highlightening plasma membranes (green arrows) in normoxia (A) and hypoxia (B). Under normoxic conditions, the plasma membrane appears smooth continuous and stable cell–cell contact, indicating high intercellular junctions and consequently maintaining high tissue integrity. In contrast, hypoxic conditions caused the cellular membrane to lose its integrity and exhibits irregularities showing less distinct intercellular junctional between adjacent cells. These changes indicate hypoxia-induced membrane leakage and junctions weakening.

### Fig. 5G

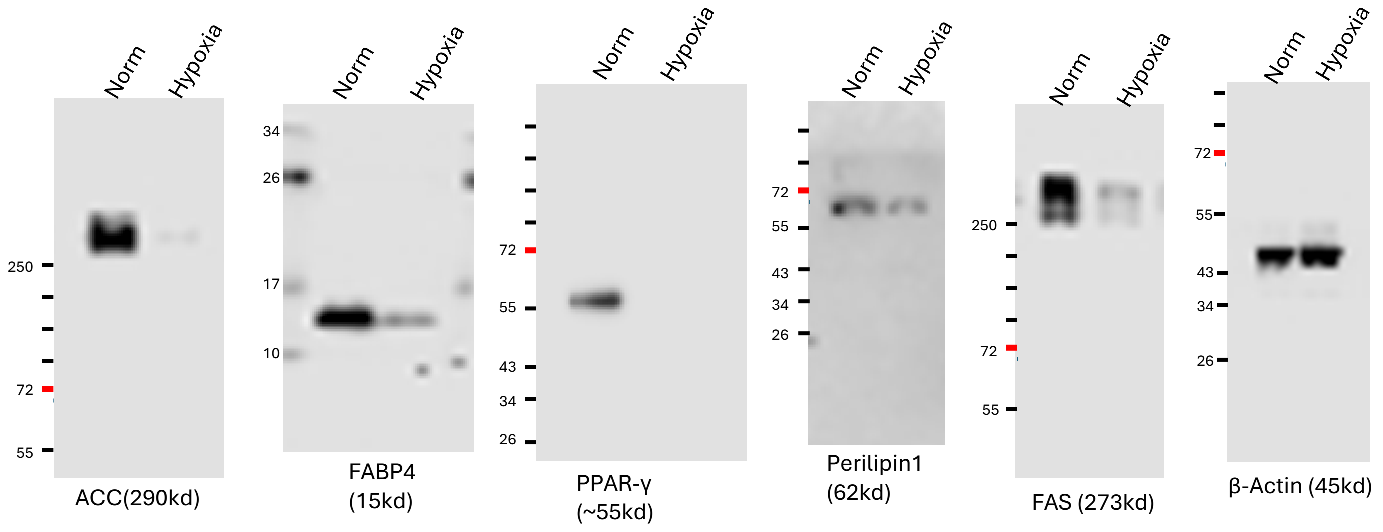


### Fig. 8A

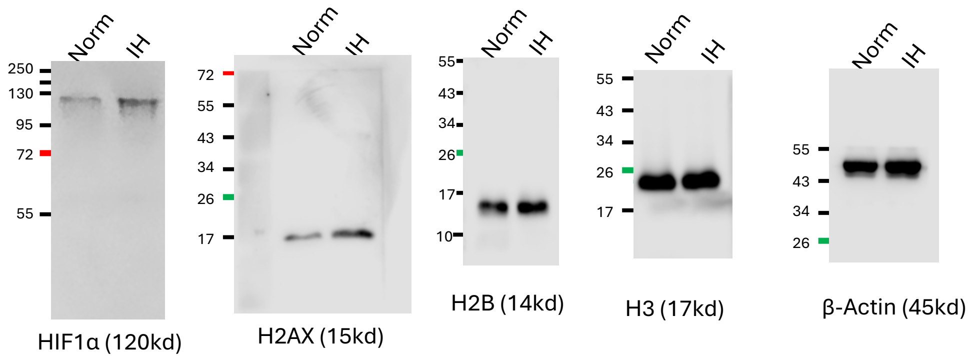


### Fig. 8E

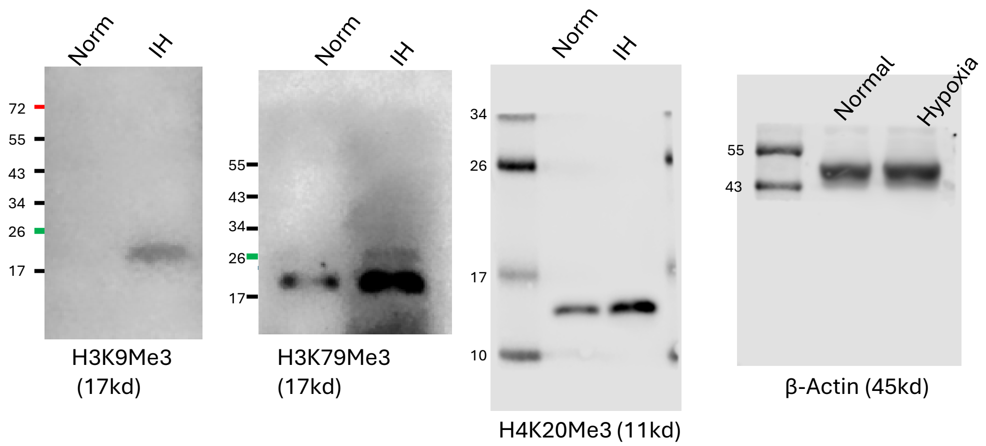


**Supplementary Fig. 10. Uncropped western blots for Fig. 5G, Fig. 8A and Fig. 8E.**

Fig. 8G

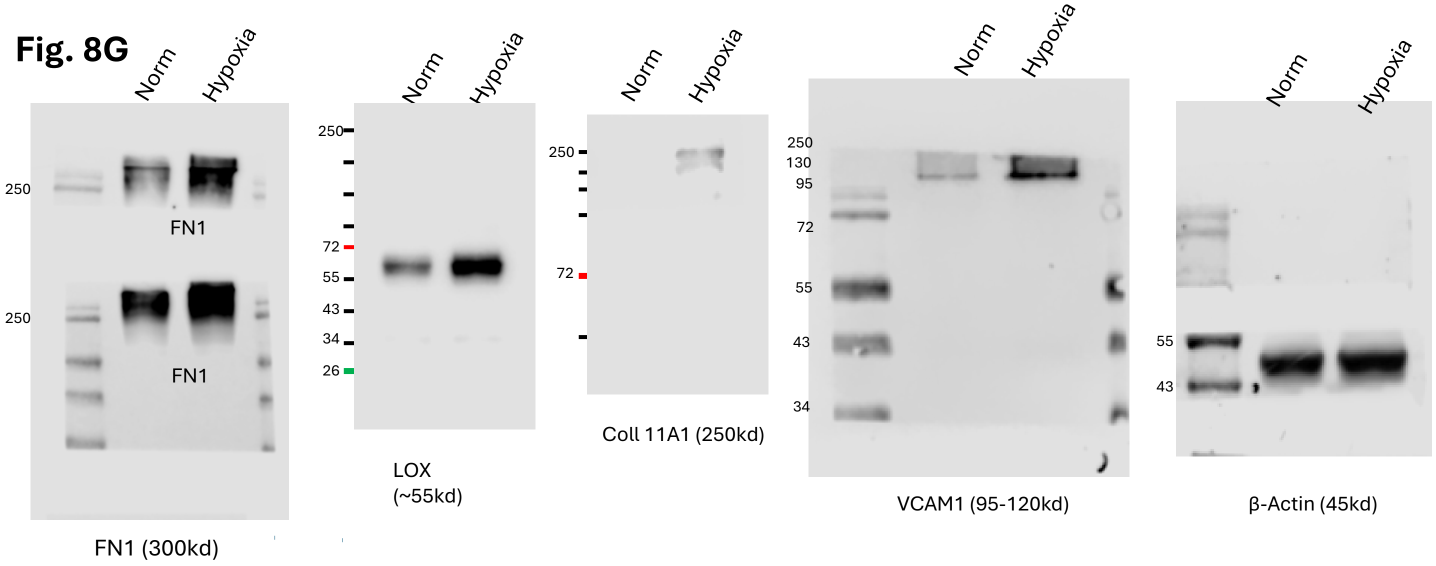


Fig. 9

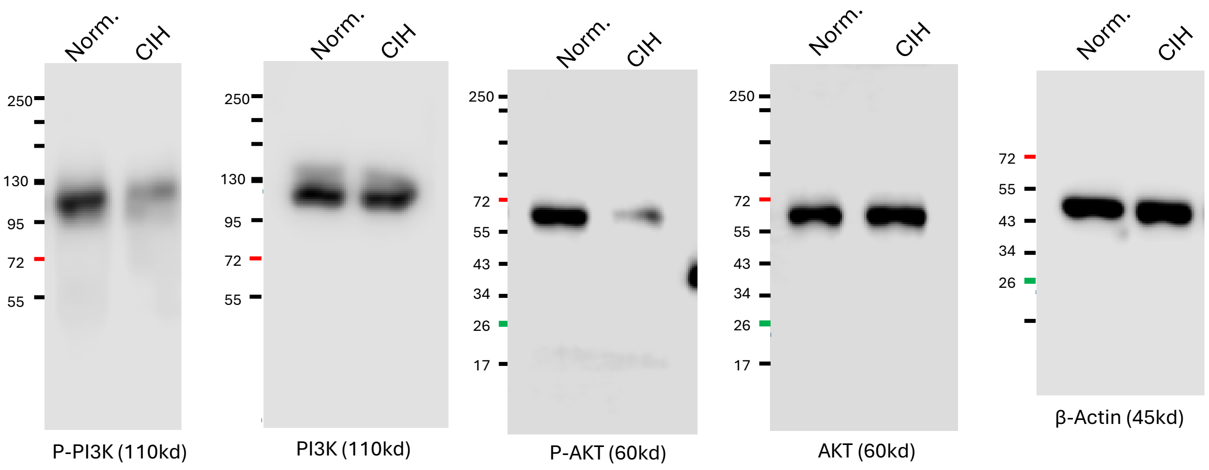


**Supplementary Fig. 10 (cont.). Uncropped western blots for Fig. 8G and Fig. 9.**
